# Mucin modulates phage infection dynamics and biofilm formation in enteropathogenic *Yersinia enterocolitica*

**DOI:** 10.64898/2026.03.24.713101

**Authors:** Sophia Goladze, Daniel de Oliveira Patricio, Ericka Allen, Reetta Penttinen, Henni Tuomala, Sheetal Patpatia, Matti Ylänne, Bent Petersen, Mikael Skurnik, Gabriel Magno de Freitas Almeida, Lotta-Riina Sundberg

## Abstract

Mucosal barriers serve as a multifunctional interface and nutrient-rich habitat for diverse microbes, including bacteria and bacteriophages. Some phages can bind to mucin glycoproteins via carbohydrate-interacting modules and provide an additional layer of mucosal immunity by shielding the underlying epithelium from invading bacteria. However, the role of mucins in shaping phage-bacterium interactions remains poorly understood. We investigated the dynamics between highly pathogenic *Yersinia enterocolitica* serotype O:8 and its mucus-adherent phage fMtkYen801 under *in vitro* mucosal environment. We assessed how mucin supplementation, varying phage doses, nutrient and temperature conditions influence phage-bacterium dynamics and biofilm formation. We found that pre-exposure to mucins led to enhanced phage replication in the bacterial host, with a 2-log increase in phage titers, and high abundance of surviving bacteria. Interestingly, mucin glycoproteins also provided *Y. enterocolitica* a nutrient source and a chemical cue to modulate its growth and biofilm biogenesis. Genomic analysis of phage-resistant bacterial variants revealed mutations in virulence, quorum sensing, and antibiotic resistance genes in both mucin enrichment and control groups, suggesting potential fitness tradeoffs during resistance evolution. Collectively, these findings highlight the importance of mucosal surfaces as an important ecological driver of phage-host interactions in *Y. enterocolitica*, a significant enteric pathogen, and emphasize the need for investigating these dynamics under complex, physiologically relevant systems to inform better phage therapy strategies against mucosal bacterial infections.

## Introduction

Mucosal surfaces represent complex and dynamic ecological interfaces for host-microbe interactions in metazoans (1–6). These interactions mainly occur through microbial adhesion to O-linked carbohydrates, which constitute a major part of mucin glycoproteins - the large macromolecular structures responsible for viscoelastic gel-forming properties of mucus (3,7). Gel-forming mucins facilitate trapping and immobilization of pathogens while also protecting the epithelium from shear stress through lubrication and hydration. Additionally, mucus enables the exchange of nutrients, oxygen and metabolites across the mucous membranes (8). The composition and thickness of mucus may influence the structure of microbiomes as well as bacterial biofilm formation, growth and virulence (7). Therefore, mucus layers are increasingly recognized not only as physical barriers against pathogen invasion and colonization but also as dynamic, responsive components of metazoan host defense and key elements of innate immunity (1,9). Moreover, the integrity of mucus and its interaction with microbiota is now acknowledged as essential for maintaining human health (7,10,11).

Alongside trillions of bacteria, archaea, fungi and viruses, phages are also major constituents of mucosal microbiota, representing 97.7% of the virome (5,7,12). In 2013, Barr et al. proposed the Bacteriophage Adherence to Mucus (BAM) model as an additional layer of non-host derived mucosal immunity. According to this model, weak binding interactions between mucin glycan residues and immunoglobulin-like (Ig-like) folds expressed on structural proteins of many tailed dsDNA phages are responsible for phage-mucin adherence (13). Combined with sub-diffusive motion of mucus-adherent phage particles into the mucosal layers, these interactions can lead to transient abundance of phages across the mucosal surfaces, thus increasing the likelihood of phage-bacteria encounters, and productive phage-mediated clearance of pathogenic bacteria (14–17). Numerous *in vitro* studies also explored the interplay between phages, their bacterial hosts and mucins as eukaryotic signals (18–24). These studies show that exposure to mucosal surfaces can modulate bacterial physiology, virulence, susceptibility to phage infection and selection for phage resistance strategies. The BAM model has been investigated in diverse model systems (15,22,23,25,26). For instance, *Flavobacterium columnare* phage FCL-2, containing Ig-like protein domain, demonstrated binding ability to primary mucus *in vivo*, leading to high phage retention on rainbow trout skin for up to 7 days after phage exposure, and protection against bacterial infection (25). Similarly, some *Pseudomonas aeruginosa* phages exhibited enhanced replication rates in the presence of mucins (23), as well as high persistence in the upper respiratory tract of mice (phage VAC3) (26). Of note, mice pretreatment with phage VAC3 before exposure to lethal *P. aeruginosa* dose was associated with higher survival rates and improved clinical outcomes compared to treatment-naïve controls (26). Similar infection preventing effect has also been found against *Vibrio cholerae* in mice and rabbits (27). Other studies in murine experimental model demonstrated that T4-like phages ΦPNJ-6 and ΦPNJ-9 can adhere to fucose residues in intestinal mucosa via Ig-like domains and reduce enterotoxigenic *Escherichia coli* colonization (28,29). Interestingly, a single dose of orally administered phage X1 treatment could also eliminate *Yersinia enterocolitica* in 33.3% of the infected mice (30), indicating the potential therapeutic value of phages during mucosal infections.

While BAM model provides valuable insights for advancing phage therapy and prophylaxis of mucosal bacterial infections, precise effects of mucin on phage-host dynamics, particularly in systems relevant to gut environment, remain poorly understood. As the efficiency of antibiotic treatments are declining due to increasing resistance among bacteria (31), filling this gap in knowledge can result in better understanding on the applicability of phage-based solutions against bacterial infections.

*Yersinia enterocolitica* is among the most commonly reported enteric pathogens in European populations (32,33), causing foodborne gastroenteritis. Antibiotic-resistant *Y. enterocolitica* strains have been isolated from various sources such as seafood (34), pork and chicken (35). Given the clinical and epidemiological significance, we investigated the impact of simulated mucosal conditions on phage replication, bacterial growth dynamics and biofilm formation using enteropathogenic *Y. enterocolitica* serotype O:8 and its newly isolated mucin-adherent phage fMtkYen801. Our findings demonstrate that exposure to mucin glycoproteins, alongside phage dosage, temperature and nutrient conditions, shape phage-bacterium dynamics. Of note, bacterial adaptation to mucosal environment prior to phage exposure led to increased phage replication and suppressed biofilm formation, suggesting that mucins act as environmental cues modulating phage infection dynamics and bacterial responses to phage challenge. These results highlight the role of mucosal environments in shaping phage-bacteria interactions, with potential implications for optimizing phage-based therapies and biocontrol strategies against mucosal bacterial infections.

## Results

### Phage fMtkYen801 is retained on mucin-coated agar plates

Phage fMtkYen801 represents a myovirus within the class of *Caudoviricetes*, lytic against *Y. enterocolitica* serotypes O:8 and O:6,30 (Supplementary Table S1). It has a 92,588 bp genome harboring 130 coding sequences (CDS), with functional annotations for 36 of them (Supplementary Table S2). Detailed phenotypic and genomic characteristics of phage fMtkYen801 are provided in Supporting Information 1. Annotated gene products include DNA metabolism, host cell lysis, and structural proteins. A tRNA Asn gene was also detected. Among the structural proteins, Pharokka identified an Ig domain-containing protein (ORF 9, 10801-11466), associated with weak interactions with glycan residues (13,14,25). Manual confirmation of the CDS using HHPred revealed high similarity to a major capsid protein (Probability: 95.84; E-value: 0.099; Score: 47.26), supporting the initial annotation, as Ig-like protein domains are frequently found on phage capsids (13,14).

The structure of Ig domain-containing protein from phage fMtkYen801 was predicted with AlphaFold2. Based on the prediction, the protein contains 7 β-strands divided into two antiparallel sheets, a hallmark of Ig constant (IgC)-type domains. This domain typically serves as a scaffold for molecular recognition and binding, which may be relevant to interactions with mucin glycans (13). Confidence metrics of the β-strands forming the Ig fold (pLDDT > 90) indicates highly reliable prediction of these regions (Fig. 1A). Electrostatic surface analysis identified positively charged patches which may facilitate interaction with negatively charged mucin glycans (Fig. 1B).

**Fig. 1.**
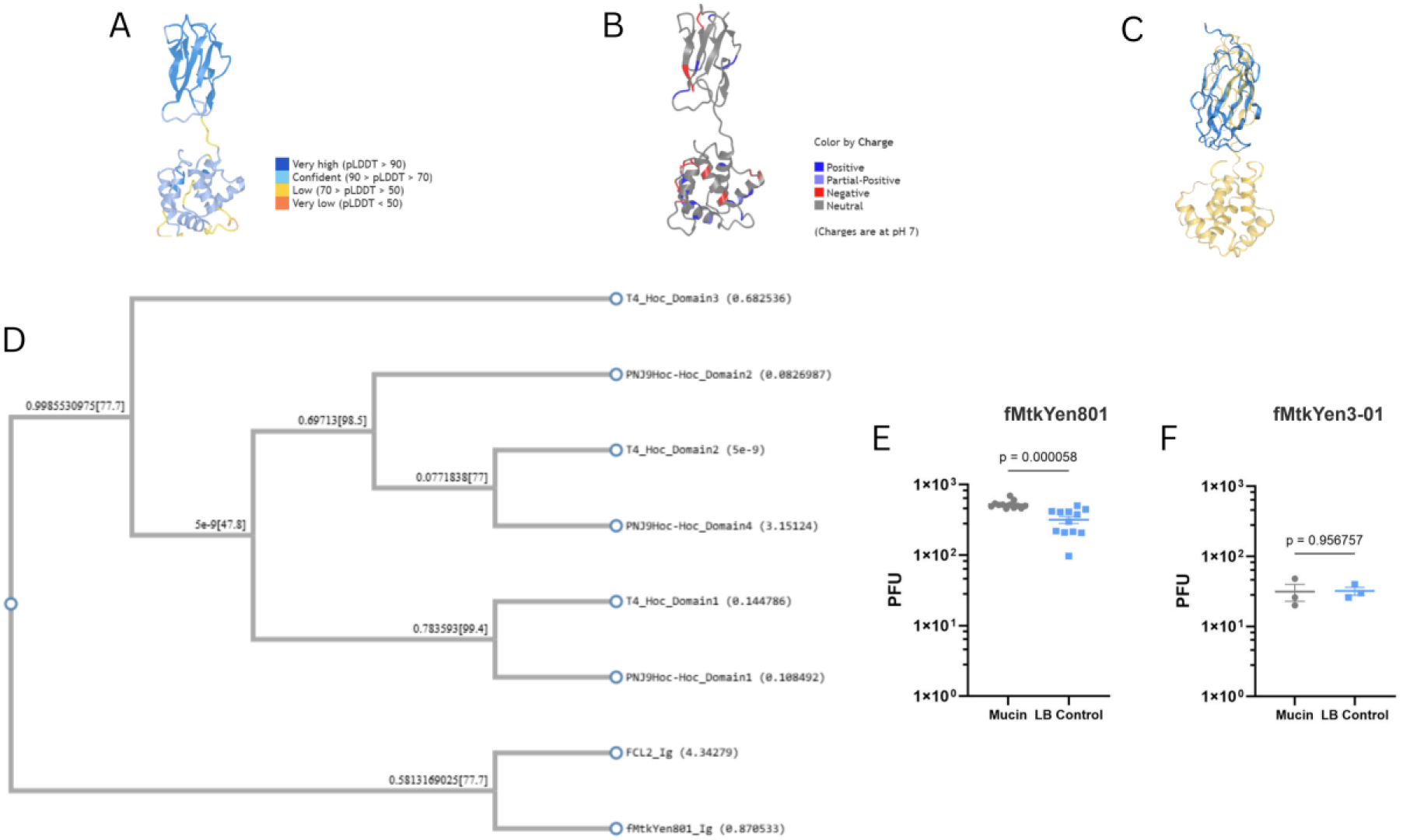
Predicted structure and mucin-binding activity of phage fMtkYen801 immunoglobulin-like (Ig-like) domain. (A) AlphaFold structural prediction of Ig domain-containing protein, illustrating the Ig-like fold (dark blue). (B) Electrostatic surface representation highlighting positively charged patches (blue) potentially mediating interactions with negatively charged mucin glycans. (C) Structural comparison between Ig-like domains of fMtkYen801 (yellow) and FCL-2 phages (blue) indicating conservation of the β-sandwich scaffold with differences in loop conformations. (D) Phylogenetic analysis of Ig-like domains between fMtkYen801, T4-like and FCL-2 phages. (E) Phage fMtkYen801 retention on mucin coated agar surfaces compared to LB control, measured by Plaque Forming Unit (PFU) counts. (F) Retention of phage fMtkYen3-01 (lacking Ig-binding domain) on mucin coated agar surfaces compared to LB control, measured by PFU counts. The error bars represent standard error of the mean (± SEM). Each treatment included 12 technical replicates for panel (E), and 3 technical replicates for panel (F). Statistical significance was assessed using unpaired two-tailed t-test.

Sequence-based phylogeny between Ig-like domains of fMtkYen801 (*Y. enterocolitica* host), FCL-2 (*F. columnare* host; ENA accession: KM873719) (25) and T4-like phages including T4 and ΦPNJ-9 (*E. coli* host; GenBank accessions NC_000866.4 and PQ100635, respectively) were analyzed using clustalw and PhyML. The phylogenetic tree showed that Ig-folds of fMtkYen801 and FCL-2 phages clustered closer together, while those from T4-like phages formed a distinct clade, highlighting the evolutionary distinctions between Ig-domains of these phage groups (Fig. 1D).

To complement the sequence-based analysis, structural comparison was performed using Foldseek. The MSA alignment confirmed a closer evolutionary distance between fMtkYen801 and FCL-2 Ig folds, compared to that of T4 phage. Specifically, the structural overlay suggested conservation of core β-sandwich scaffold between Ig folds of fMtkYen801 and FCL-2 phages. Notably, the alignment also identified divergence between the orientation and length of loop conformations, which often facilitate ligand recognition and may be relevant to glycan binding (Fig. 1C). For comparison, T4 phage fold was shown to be closer to ΦPNJ-9 (14,25,29).

To confirm *in silico* prediction, we investigated phage binding ability to the purified porcine gastric mucin (PGM) *in vitro*. A phage suspension (2.5×10^3^ PFU/mL) was applied to LB agar surfaces with or without 1% (w/v) mucin coating. After 30 min incubation, excessive liquid was removed from the plates, and phage retention to the agar surfaces was estimated using double-overlay agar assay. As an experimental control, we performed an independent assay using phage fMtkYen3-01 (2.5×10^2^ PFU/mL) and its host, *Y.enterocolitica* serotype O:3 (6471/76-c) (36). In contrast to fMtkYen801, this phage does not encode Ig-like domain, and therefore, served as a negative control for the mucin-binding assay. Phage fMtkYen801 was preferentially retained in mucin-containing plates compared to standard nutrient media *(p=*0.000058), with 1.64-fold increase in adherence to mucin coated agar surface, while fMtkYen3-01 did not show significant difference between mucin and LB treatments (Fig. 1E).

### Mucin modulates post-infection growth dynamics in *Y. enterocolitica* serotype O:8 in an MOI-dependent manner

To investigate the effect of mucin glycoproteins on interactions between a mucin binding phage and its bacterial host, *Y. enterocolitica* O:8 was infected with fMtkYen801 at Multiplicities of Infection (MOIs) of 0.1, 0.01 and 0.001, and incubated at 25°C for 60 h, under varying mucin concentrations (0-0.2% [w/v]). Growth dynamics monitored by optical density (OD_600_) revealed that phage suppressed bacterial growth in all tested conditions, and the subsequent recovery phase was jointly affected by mucin supplementation and initial phage dose. At MOI 0.1, mucin-supplemented cultures initiated growth recovery at 36 h post-infection, while regrowth in standard nutrient medium began only after 48 h. At 60 h post-infection, final OD_600_ values were significantly higher in all mucin treatment groups compared to no-mucin control (*p* < 0.0001) (Fig. 2A). In contrast, at lower MOIs only limited increase in OD_600_ was observed, leading to significantly lower end-point values in 0.2% mucin treatments compared to the standard nutrient medium (*p* = 0.0006 for MOI=0.01, and *p* < 0.0001 for MOI = 0.001) (Fig. 2B-C). Interestingly, mucin supplementation alone did not significantly alter growth dynamics of control bacterial cultures (*p* > 0.05) (Fig. 2D). Furthermore, phage abundance measured in 60 h supernatants remained unaffected by mucin enrichment under the tested conditions (*p* > 0.05) (Fig. 2E-G), indicating that observed differences in post-infection growth dynamics in *Y. enterocolitica* O:8 are unlikely to be related with altered phage replication.

**Fig. 2.**
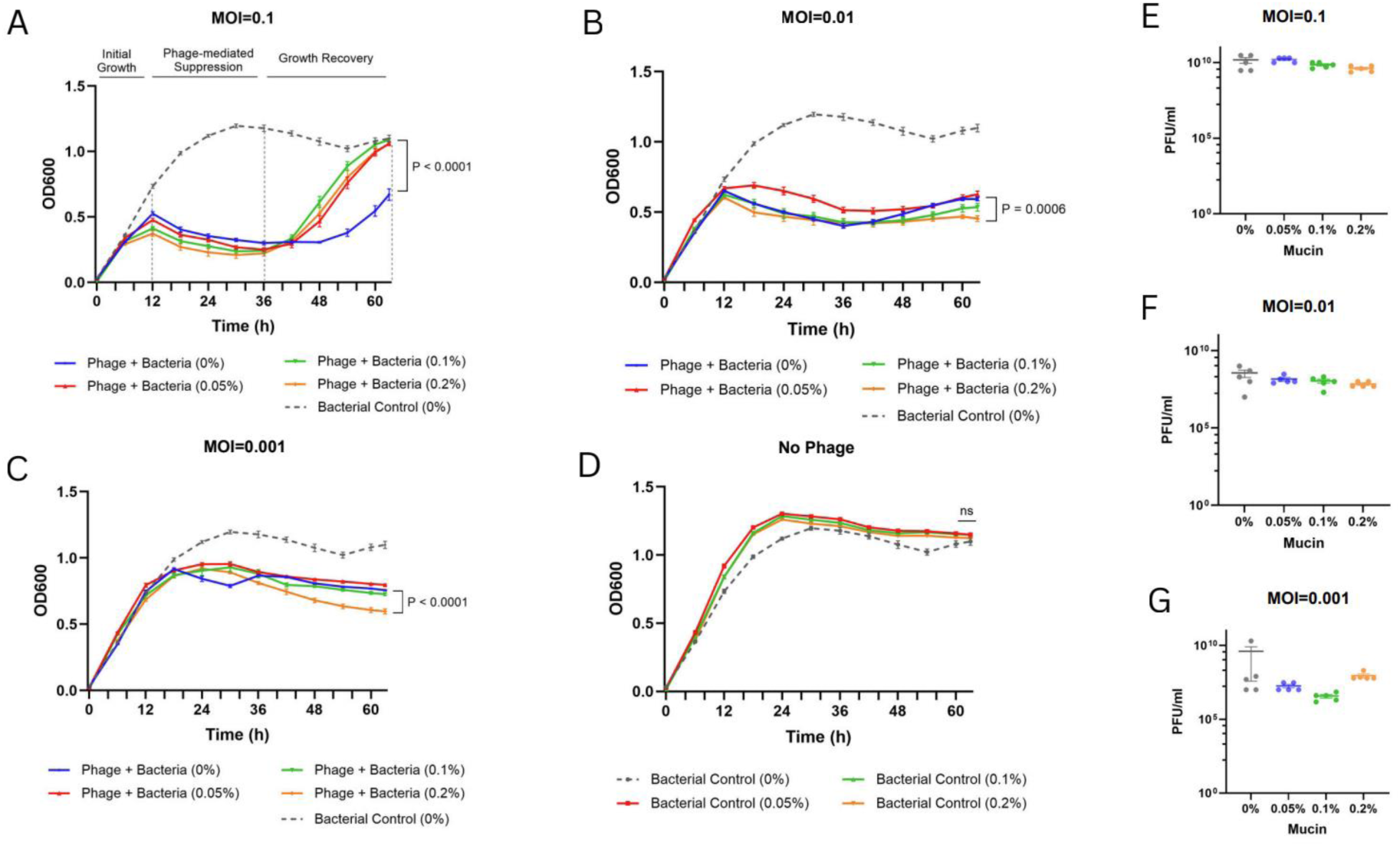
Bacterial growth dynamics and phage abundance in *Y. enterocolitica* O:8 under varying phage-to-bacteria ratio (MOI) and mucin concentrations. (A-C) Bacterial cell density dynamics (OD_600_) at MOI of 0.1, 0.01, or 0.001. (D) Bacterial growth without phage infection. Data are shown as mean ± SEM (n = 10 technical replicates per condition). (E-G) Phage abundance (PFU/mL) in culture supernatants at 60 h post-infection at MOI of 0.1, 0.01, or 0.001. Data are shown as mean ± SEM (n = 5). Statistical significance was assessed using two-way ANOVA with Tukey’s multiple-comparison test for panels A-D, and one-way ANOVA with Tukey’s post hoc test for panels E-G.

The MOI-dependent effect of mucin on post-infection growth dynamics was reproduced in an independent assay performed under the same conditions as in Fig. 2. Namely, mucin-supplemented cultures showed consistently lower cell densities compared to no-mucin controls at low MOIs (0.01 and 0.001; *p* < 0.001 and *p* < 0.0001, respectively), higher recovery at MOI of 0.1 (p < 0.0001), and no significant difference at MOI of 1 between mucin and no-mucin treatments (*p* = 0.3532) (Fig. S6). To verify these findings in an additional *Yersinia* model, we performed an independent 60 h time-course infection assay in *Y. enterocolitica* O:3 and a non-mucin binding phage, fMtkYen3-01. In this system, mucin supplementation did not exhibit consistent effects on bacterial growth dynamics under varying MOIs (Fig. S7). Interestingly, bacterial doubling time, calculated from the exponential growth phase in both co-culture experiments (Fig. 2, Fig. S7), was not significantly affected by mucin exposure alone (Fig. S9), suggesting a possible link between mucin-associated phenotype previously observed in *Y. enterocolitica* O:8 and phage fMtkYen801-specific properties.

### Bacterial pre-adaptation with mucin influences phage-bacterium dynamics under varying environmental conditions

To dissect the effects of mucin on phage-host dynamics under varying environmental conditions, fMtkYen801 and its host were co-cultured at MOI of 0.1 with or without mucin supplementation, and incubated at 25°C, or 37°C for 24 h, under constant agitation. In addition, the experiment included treatment groups where phage or bacteria were adapted to mucin for 2 h prior to infection. Bacterial pre-adaptation with mucin significantly increased viable cell counts (CFU/mL), as well as phage replication (PFU/mL), compared to simultaneously co-cultured, or phage pretreatment groups (Fig. 4A-B). In comparison, mucin pre-adaptation of *Y. enterocolitica* O:3 did not affect replication of the non-mucin binding phage fMtkYen3-01 (Fig. S8), indicating that these effects are linked to specific phage-host system.

**Fig. 4.**
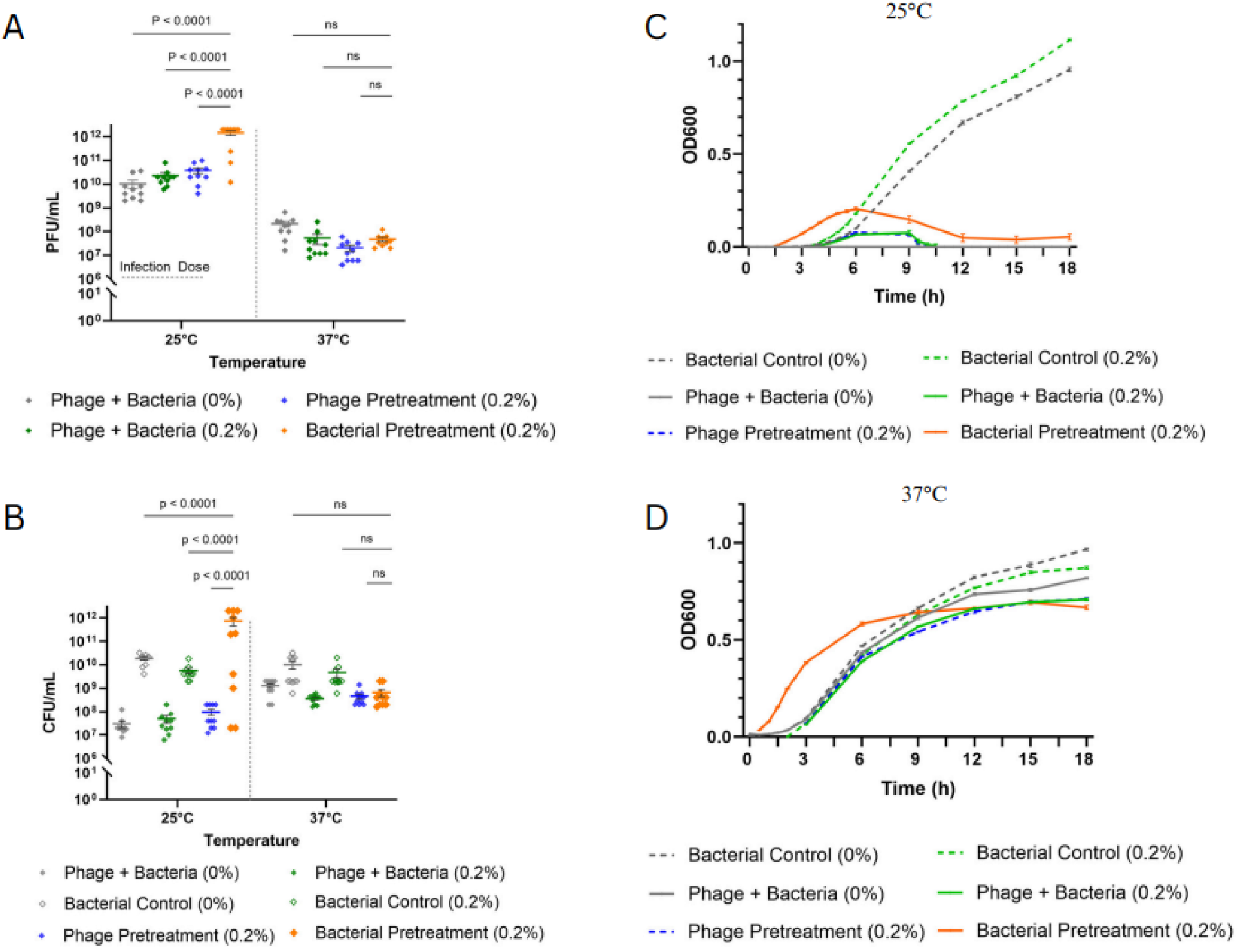
Bacterial growth and phage replication under varying mucin supplementation and temperature conditions. (A-B) Phage replication and surviving bacterial cell counts (C) Bacterial growth curve at 25°C (D) Bacterial growth curve at 37°C. Error bars indicate mean ± SEM (n = 10 technical replicates). Statistical significance was assessed using two-way ANOVA followed by Tukey’s multiple comparisons tests.

Phage infection efficiency was significantly reduced at 37°C, compared to 25°C (Fig. 4), possibly, due to decreased availability of receptors leading to reduced phage adsorption.

This hypothesis was further supported by screening the engineered LPS-mutant strains on phage susceptibility, indicating that fMtkYen801 uses O antigen (O-ag) as one of the receptors for adsorption (Fig. S2). Moreover, O-side chain of LPS is known to be expressed only at low temperatures in *Y. enterocolitica* (≤ 25°C) (37–39).

Next, we examined the effects of mucin on phage-host interactions under varying nutrient conditions. Mucin-adapted and non-adapted bacterial cultures were infected with fMtkYen801 at MOI of 0.1, and growth dynamics monitored by OD_600_ for 18 h in nutrient-rich (LB) or nutrient-deficient (dH₂O) conditions. Interestingly, in nutrient-deprived conditions, phage-infected cultures showed higher cell densities than non-infected controls (*p* < 0.0001), while pre-exposure to mucin led to further increase in OD_600_ values (Fig. 5B). This pattern was not observed in nutrient-rich conditions (Fig. 5A).

**Fig. 5.**
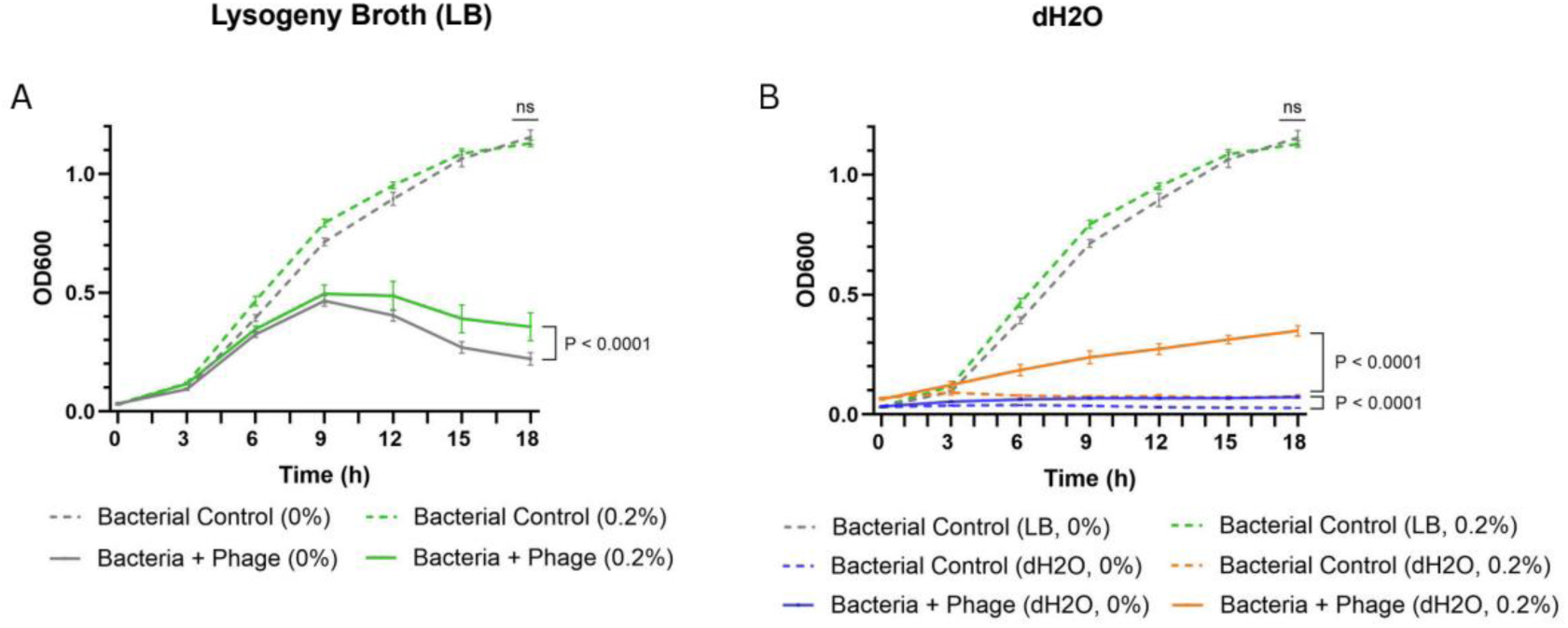
Bacterial growth with and without pre-adaptation to mucin under nutrient-rich and -deprived conditions. (A) Growth in standard nutrient media (LB) with or without mucin exposure (B) Growth in dH₂O with or without mucin exposure. Error bars represent mean ± SEM (n = 10 technical replicates). Data was analyzed using two-way ANOVA, followed by Tukey’s multiple comparison test.

### Environmental factors, including mucin exposure, modulate biofilm formation in *Y. enterocolitica*

To investigate the influence of environmental factors on biofilm development in *Y. enterocolitica* O:8, we quantified surface-attached biofilm biomass using crystal violet (CV) staining under mucin supplemented conditions (0%, 0.05%, 0.1% or 0.2%) after 48 h static incubation. All mucin treatment groups showed reduction in biofilm biomass compared to no-mucin control (Fig. 6A).

**Fig. 6.**
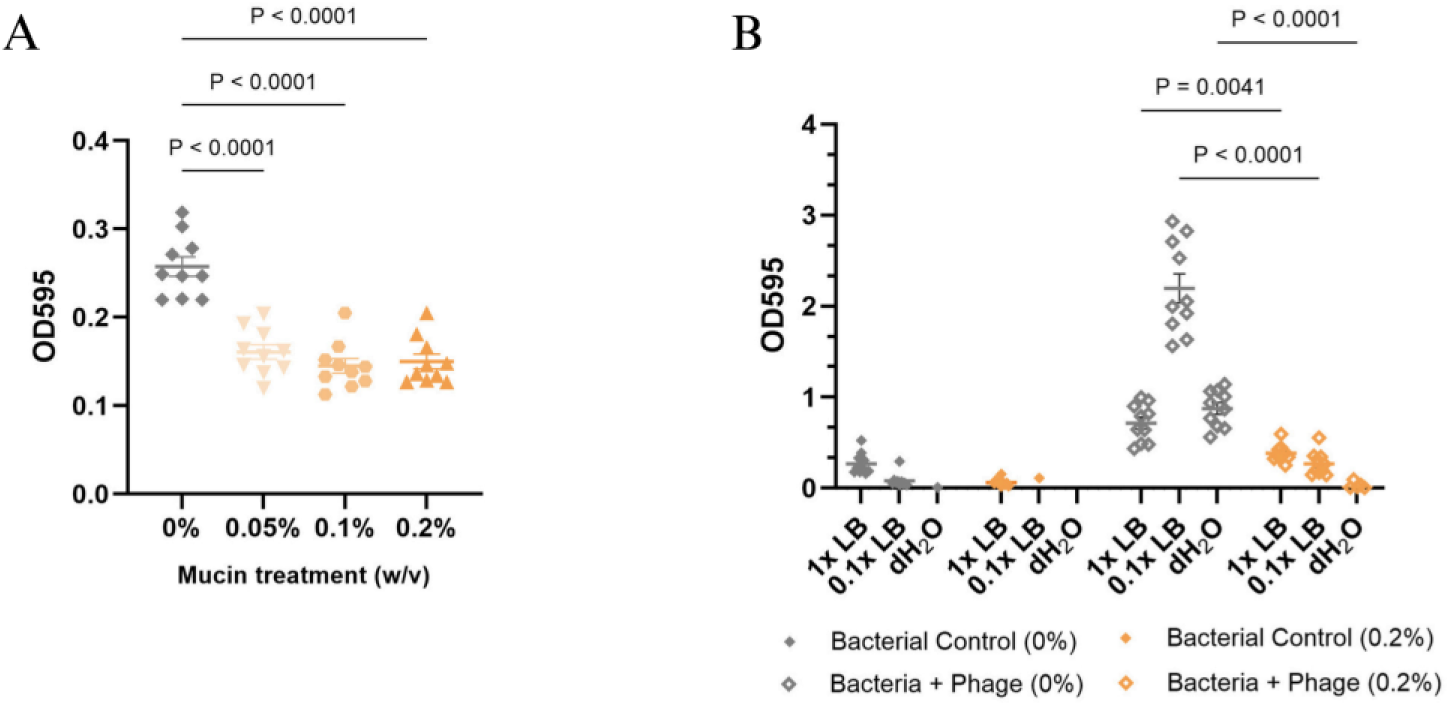
Effects of nutrient availability, mucin supplementation, and phage infection on biofilm formation in *Y. enterocolitica* O:8, quantified using crystal violet (CV) assay. (A) Surface-attached biofilm biomass in varying mucin concentrations. (B) Biofilm formation under varying nutrient conditions, mucin supplementation, or phage exposure. OD_595_ values are background-corrected using medium-only wells, with or without mucin. n = 10 technical replicates per condition. Error bars represent standard error of the means (± SEM). *p* values are indicated in the figure. Statistical significance was calculated using one-way (A) and two-way (B) ANOVA with Tukey’s post hoc test.

Next, we challenged bacteria with phage at MOI of 0.1 and exposed them with varying nutrient conditions (1x LB, 0.1x LB, dH₂O), with or without mucin supplementation. *Y. enterocolitica* showed minimal CV signal under all tested conditions. However, phage infection led to significant increase in biofilm formation, especially under nutrient limitation, raising mean OD_595_ values from 0.08 to 2.196 in 0.1x LB (*p* <0.001). As observed previously (Fig. 6A), mucin supplementation reduced the amount of biofilm in all tested nutrient conditions, keeping the mean values near the baseline levels (*p* <0.001) (Fig. 6B). Cell-free wells, containing sterile medium with or without mucin enrichment, showed low background absorbance (≤ 0.2) (Fig. S10A). Notably, differences in the biofilm biomass were independent from planktonic cell densities measured after 48 h static incubation (Fig. S10B).

Building on these findings, we compared biofilm formation between the ancestral *Y. enterocolitica* (wild-type) and Bacteriophage Insensitive Mutant (BIM) strains (efficiency of plating (EOP) of 0), derived from a 60 h co-culture experiment in the presence (BIMs 1-3) or absence (BIMs 4-6) of 0.2% (w/v) mucin. Interestingly, most of the tested BIMs exhibited significantly higher levels of biofilm biomass compared to the ancestral strain (Fig. 7A). Consistent with the previous experiments, mucin reduced the amount of adhered biofilm biomass in all tested strains (*p* <0.0001) (Fig. 7B).

**Fig. 7.**
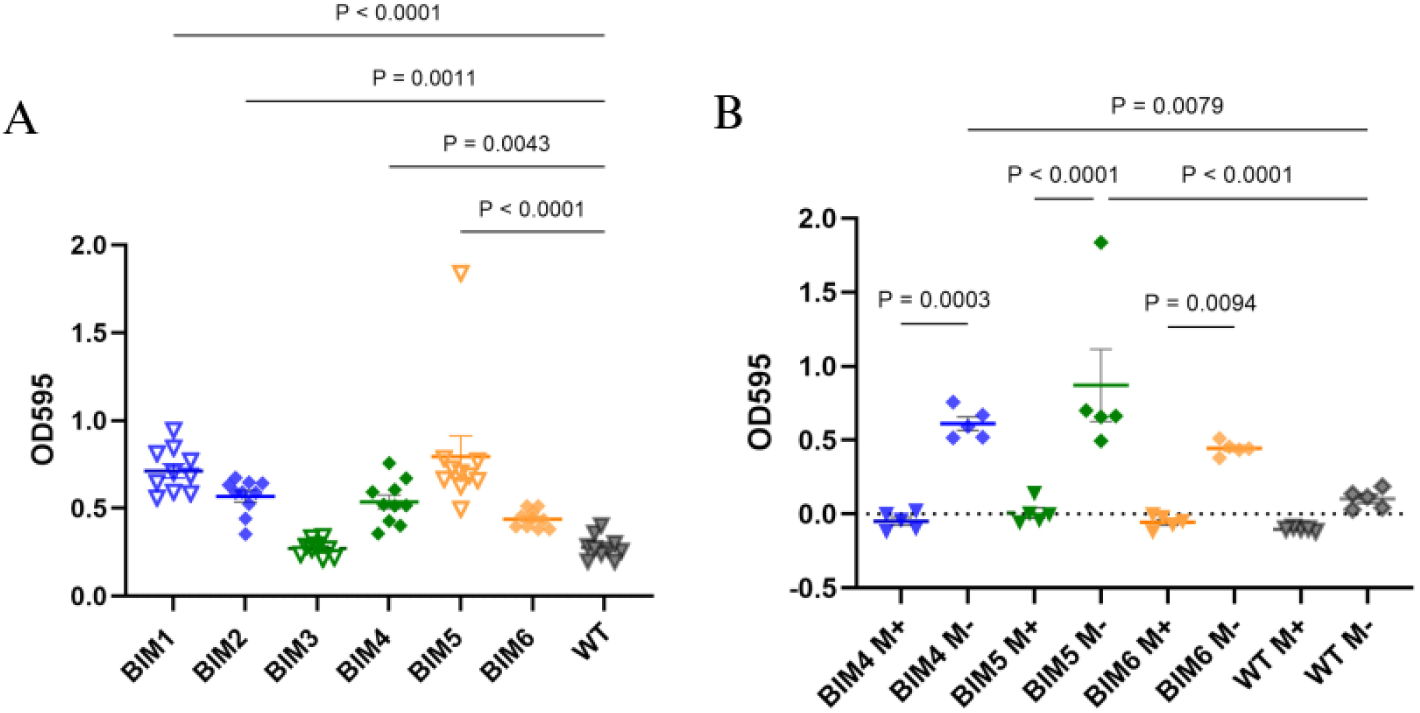
Biofilm formation in the ancestral *Y. enterocolitica* and Bacteriophage Insensitive Mutant (BIM) strains. (A) BIMs show increased adhered biofilm biomass compared to the ancestral strain (n = 10 technical replicates per group). (B) Mucin reduces biofilm formation in both ancestral strain and BIMs (n = 5). OD_595_ values were background-corrected using medium-only wells, with or without mucin. Error bars indicate mean ± SEM. Statistical significance was assessed using one-way ANOVA with Dunnett’s (A) and Tukey’s (B) post hoc tests. Exact *p* values are provided in the figure.

### Comparative genomic analysis of phage resistant mutants

Comparative genomic analysis was performed on the ancestral *Y. enterocolitica* strain, and single BIM isolates generated under mucin-supplemented or standard nutrient conditions. The ancestral strain was used as a reference for identifying mutations associated with phage resistance phenotypes in BIMs. The quality-filtered BIM reads were assembled using Flye, while the draft assembly of the reference genome was further improved by polishing with DNBseq reads, producing the largest contig with 4,587,823 bp, GC content of 47.24%, and N50 of 4,582,530 bp. Genome completeness evaluated using Busco yielded a score of 99.2%, 97.6%, 99.2% for BIM from standard nutrient medium (BIM 1), BIM from mucin treatment group (BIM 6) and the ancestral strain, respectively, indicating a high-quality assembly.

Genome annotation using Prokka identified 4119 CDSs, 22 rRNAs, 85 tRNAs and 1 tmRNA in the wild-type strain (Supplementary Tables 3-5). Slight variation was observed in gene content of BIMs compared to the ancestral genome, with 4150 CDSs, 4258 genes in 0.2% mucin treatment, and 4152 CDSs and 4260 genes in standard nutrient conditions.

The whole genome alignments using BRIG and ProgressiveMauve did not reveal any major structural mutations between the ancestral and BIM strains (Fig. S11). Padloc v2.0.0 detected conserved phage defense systems, including type II restriction modification, SoFic, and AbiE in both, ancestral and BIM genomes (Supplementary Table 6). PHASTEST identified seven prophage regions in ancestral, and six in BIM genomes, however, only three prophages in each strain were annotated as intact (score > 90) (Fig. S12; Supplementary Tables 7a-7c).

To identify mutations in BIM genomes relative to the ancestral strain, variant calling was performed. Comparative genomics identified 102 mutations across BIM genomes, which were further filtered to include only high and moderate impact polymorphisms. Kyoto Encyclopedia of Genes and Genomes (KEGG) pathway mapping and annotation of the mutated genes enabled identification of the gene pathways affected by phage resistance in both treatment groups (Supplementary Tables 7a-7c). Here, we focus on genes responsible for metabolism, virulence, quorum sensing, and antimicrobial resistance. Majority of mutations (95%) were shared between BIMs from 0% and 0.2% mucin treatment groups (Fig. S13), including a single base deletion identified in *mrcB* gene (contig_1: g.3,755,152del), encoding Penicillin-binding protein 1B, and responsible for peptidoglycan biosynthesis (40). Interestingly, two mutations were detected in *pdeR* gene (Cyclic di-GMP phosphodiesterase PdeR), involved in quorum sensing (41). The gene carried 4 bp deletion at position 4,539,163 (contig_1: g.4539163-453966del), and 3 bp deletion at 4,539,729 (contig_1:g.4539729-4539731del) with predicted frameshift mutation (p.Gly90fs), and conservative in-frame deletion (p.Lys278del). Both, *mrcB* and *pdeR* share a demonstrated link to biofilm formation (42,43). All phage resistant isolates also carried deletions in two genes encoding hypothetical proteins (at positions 1,671,635 and 919,444, respectively), predicted to be involved in secretion system and biofilm biogenesis based on KEGG annotation.

High-impact variants were detected in genes potentially involved in two-component signal transduction systems and bacterial motility. Specifically, *cheB* gene, encoding Protein-glutamate methylesterase/protein-glutamine glutaminase, showed a 3 bp deletion (contig_1:g.1822339-1822341del), leading to disruptive in-frame deletion (p.Gly167del), while *tsr_4*, encoding Methyl-accepting chemotaxis protein I, harbored 2 bp deletion (g.4123059_4123060del), resulting in frameshift and stop codon loss (p.Ter527fs). Frameshift deletion was also identified in the predicted two component system gene, *degP*, which is additionally linked to cationic antimicrobial peptide (CAMP) resistance. Lastly, shared polymorphisms by BIMs included a single nucleotide deletion (contig_1:g.1017292-1017293del) in *metl_2* gene, encoding D-methionine transport system permease protein MetI, with predicted frameshift mutation at amino acid 166 (p.Ala166fs). Similarly, genes encoding Glutamate/aspartate import permease protein GltJ, L-cystine transport system permease protein YecS, Vitamin B12 import ATP-binding protein BtuD and Iron(3+)-hydroxamate import ATP-binding protein FhuC, responsible for signaling and cellular processes, carry 1-2 bp deletions, likely resulting in truncated protein and functional disruptions (p.Pro171fs, p.Met30fs, p.Glu480fs and p.His253fs, respectively). All the above-noted genes belong to ABC transporters, facilitating vital nutrient acquisition, bacterial pathogenesis and virulence (44).

Interestingly, phage-resistant isolate from mucin enriched conditions revealed a unique nonsense mutation in *manC_1* (Mannose-1-phosphate guanylyltransferase 1), involved in GDP-mannose biosynthesis, crucial process in cell surface polysaccharide metabolism (45). The mutation resulted in a premature stop codon at amino acid position 71 (p.Gln71*). A unique missense mutation (p.Gln241Leu) was also identified in gene *gmd* (GDP-mannose 4,6-dehydratase) of a BIM isolate from standard nutrient conditions. *gmd* is involved in nucleotide sugar biosynthesis and plays an essential role in GDP-fucose production (46). Both, *manC* and *gmd* represent gene clusters of O-antigen part of LPS, directly linked to resistance to phages, cationic antimicrobial peptides, and bacterial virulence (37,47,48). These results suggest that mutations in O-antigen genes are primarily driven by phage-mediated selection pressure rather than environmental factors, such as the presence of mucin.

## Discussion

Mucosal surfaces are complex ecosystems that harbor a high diversity of microbial communities, including commensal and pathogenic bacteria and bacteriophages (6).

Our study was motivated by Bacteriophage adherence to mucus (BAM) model, proposed by J. Barr et al. over a decade ago, suggesting that phages are capable of adhering to mucin O-glycans, providing non-host-derived, innate immunity against bacterial pathogens (14,21,26). Ig-like domains, expressed on highly immunogenic outer capsid protein Hoc, together with other phage-encoded mucin-interacting proteins, are considered central in phage adhesion and retention in the host mucosal environment (12,14,49,50). It is noteworthy that Ig-like domains are encoded by approximately 25% of tailed dsDNA phages (15,51), many of which may be naturally interacting with carbohydrates in their ecological niches. Additionally, Green et al. showed that some phages possess ability to bind to mammalian glycans through tail fiber proteins, and by ensuring more frequent encounter with bacterial hosts, enhance infectivity in mucin-rich environments (21). So far, BAM model dynamics have been demonstrated in various systems, including *E. coli*, *P. aeruginosa*, *F. columnare and Vibrio anguillarum,* among others (15,22,23,25,26). Despite the importance of mucosa, the impact of its main component, mucin glycoproteins, on the outcomes of phage-host interactions is not fully understood in many bacterial species (14,19,25,52). To extend this concept, we investigated properties relevant to BAM model using a newly isolated *Yersinia* phage fMtkYen801. fMtkYen801 revealed 1.64-fold higher *in vitro* binding ability to porcine gastric mucin, compared to standard nutrient conditions, potentially through Ig-like domain. Although BAM model has been previously associated with increased local phage concentrations, we did not see the direct link between phage retention on mucus-enriched agar surfaces and enhanced phage replication relative to standard nutrient conditions, evidencing that these are unlinked phenotypes and in line with other phages tested on this sense (23,25). Nevertheless, we speculate that phage adherence to mucus layers, as a key aspect of the BAM model, together with the ability to infect the host under mucosal conditions, can facilitate preventive protection against bacterial infections (53).

In contrast to the above-mentioned results, we discovered that bacterial adaptation to mucins prior to phage infection led to significant increase in phage replication, together with high abundance of surviving bacteria. Moreover, mucin treatment groups showed reduced surface-attached biofilm biomass and favoured planktonic growth in *Y. enterocolitica* O:8 model. These findings are consistent with the studies by Almeida et al. (2019; 2024), which demonstrate that bacterial exposure to mucins can enhance their susceptibility to phage infection, while modulating bacterial virulence (23,25). Additional studies further support these findings. For instance, mucin mediated enhancement of *Streptococcus mutans* growth led to four-log increase in phage concentration (18). Furthermore, improvement in phage infection efficiency was observed during bacterial host exposure to human heparan sulfated proteoglycans, as well as in mucus-producing human epithelial HT29 cell line (21,54). One mechanistic explanation for this phenotype may be increasingly recognized glycan-mediated modulation of pathogen physiology, including global transcriptional changes, virulence, growth, and potentially, phage receptor expression, enabling successful virus adsorption and predation. However, this effect seems to be highly species and context dependent. For instance, mucin has been shown to upregulate virulence in *Acinetobacter baumannii* (4). In contrast, mucin glycans downregulate virulence gene expression, and demonstrate biofilm dispersal effect in *Streptococcus pneumoniae* and *P. aeruginosa* PAO1 cultured in HT29 cells, as well as in relevant animal models (55,56). This variability emphasizes the importance of studying interactions between mucins, microbes and the metazoan host in physiologically relevant, complex environments.

For successful colonization of gastrointestinal (GI) tract, bacteria need to adapt to highly complex environmental conditions. In 2023, Carroll-Portillo et al. showed that efficacy of phage infection in luminal environment was not solely defined by applied phage dosage, but was also significantly affected by external factors, such as mucin and agitation in the microenvironment (52). Similarly, our results suggest that while mucin and phage dosage strongly influence phage infection dynamics, post-infection cell recovery and biofilm formation in the bacterial host, additional environmental factors, such as nutrient availability, and temperature may further alter these interactions. Specifically, we observed the ability of the host bacteria to grow in mucin-enriched sterile pure water. Many bacterial species can adhere to mucin glycoproteins via lectin-type adhesins and degrade glycans to obtain the necessary nutrient resources, supporting their growth (5,57–59). Temperature-dependent growth enhancement in mucin supplemented medium has also been reported for *Y. enterocolitica* (60); however, we saw the growth phenotype only under nutrient deprived conditions, and exclusively in the presence of phage. One possible explanation is that initial low-level phage infection in nutrient-limited conditions induced bacterial cell lysis, releasing intracellular nutrients into the environment, that subsequently supported the growth of phage-resistant bacterial subpopulations (61). Alternatively, we speculate that phages may possess carbohydrate-active enzymes capable of modifying mucin glycans or their protein backbones, thereby facilitating bacterial access to glycan-derived nutrients (4,12,62). It is noteworthy that phage genome annotation did not identify any known enzymes directly linked with mucin glycan degradation, however, 72% of the predicted genes encode hypothetical proteins which could potentially harbor uncharacterized enzymatic functions. Validation of these hypotheses would require further experimental investigation. In 2021, Carroll-Portillo et al. suggested that combination of phage therapy and prebiotics could improve therapeutic outcomes by providing sufficient amount of short chain fatty acids (SCFAs), essential substrates for intestinal epithelial cells (49,63). Building on the observation that *Y. enterocolitica* can use PGM glycans as a nutrient source under phage exposure, we further speculate that prebiotic and probiotic supplementation of GI tract could act synergistically with phage therapy by stimulating thickness of mucosal barrier, increasing abundance of beneficial mucin degrading bacteria and their metabolites, alleviating competition between pathogenic and commensal bacteria for native mucin glycans, and thereby, preventing the damage and inflammation of mucosal barrier. However, to the best of our knowledge, direct evidence for successful synergy between phages and pro- or prebiotics remains lacking. It should also be noted that protein backbone and O-glycans of commercially available PGM differ from human colonic mucins, potentially limiting direct translational relevance of our findings (64,65). However, when compared to primary mucus, PGM had similar effects for mucosal-associated phenotypes (25), and is so far the most common mucin source used for BAM model studies (14,15,25,26).

Interestingly, phage fMtkYen801 infectivity was significantly reduced at 37°C both in standard nutrient medium and simplified mucosal microenvironment, likely due to downregulation of O-ag gene expression at higher temperatures (39,66). This hypothesis was supported by phage susceptibility testing of engineered LPS-mutant *Y. enterocolitica* strains, as O-ag-deficient mutants exhibited resistant phenotype, indicating that fMtkYen801 uses O-ag as one of the receptors for adsorption (67,68). Additionally, BIM genome analysis confirmed that resistance was associated with nonsense mutations in O-ag gene clusters (*manC_1*, *gmd*). Alternatively, temperature might influence bacterial physiology, or mucin properties, leading to reduced phage replication, however it has not been directly assessed in this study. Overall, these results highlight that within luminal microenvironment of natural infection models, pathogen exposure to mucins, adaptation to body temperature (37°C) and competition for nutrient resources can influence phage-bacteria interactions and affect phage therapy outcomes.

Mucins have been shown to increase frequency of resistance mutations to protect the bacterial populations from phage infections on the mucosal surfaces (19,24). For better understanding of the selection for phage resistance in *Y. enterocolitica*, we analyzed whole genome sequences of single BIM isolates from mucin enriched or standard nutrient conditions. We observed multiple genomic changes, likely leading to trade-offs between phage resistance, biochemical properties and virulence. Most of the mutations were shared across BIMs, e.g. polymorphisms in virulence genes, including *pdeR,* responsible for quorum sensing and biofilm biogenesis, as well as in several hypothetical proteins predicted to be involved in secretion systems. This finding aligns with our experimental results, showing significantly enhanced capacity for biofilm formation in phage resistant mutant strains. Additionally, we found mutations in genes responsible for cationic antibiotic resistance, suggesting the possible changes in antibiotic susceptibility of phage resistant variants. However, in this study, this aspect was not further investigated. High impact deletions were identified in genes associated with two-component system (*degP*), motility and chemotaxis (*cheB, tsr_4*), and ABC transporter gene cluster, critical for nutrient uptake and virulence (*gltJ, yecS, btuD, fhuC, metI*). Interestingly, nonsense mutations were found in O-ag genes (*manC_1, gmd*), known as one of the virulence factors for *Y. enterocolitica* and main recognition receptors for *Yersinia* phages (69). These mutations likely resulted in truncated or dysfunctional proteins, thereby inhibiting phage infection at the initial stage of adsorption. These findings position phage selection pressure, rather than the mucosal environment as the main driver of mutations associated with phage resistance, virulence and metabolism in the bacterial host.

Collectively, consistent with previous research, observed genotypic and phenotypic patterns highlight dynamic and context-dependent nature of phage-bacteria interplay in mucosal environments. Further studies in physiologically relevant environments that reflect complexity of human biology could advance understanding of mucosal microbial ecology and contribute to better phage therapy outcomes.

## Materials and methods

### Bacterial strains and culture conditions

Bacterial strains used in this work are listed in Supplementary Table 1. *Yersinia enterocolitica* strain 8081-c serotype O:8 (1) was used as phage isolation host. All *Yersinia* strains were cultured at room temperature (RT) ranged between 20 and 25 °C, in Lysogeny Broth (LB) (10 g/L Tryptone, 5 g/L Yeast Extract, 10 g/L NaCl, pH 7.0). When needed, LB was supplemented with 0.4% or 1.5% (w/v) agar. Simulated mucosal conditions were generated using purified porcine gastric mucin (Sigma-Aldrich, catalog no. M1778). A 2% (w/v) stock solution was prepared in sterile water, autoclaved and diluted in respective nutrient media to obtain the desired mucin concentrations.

### *In silico* prediction and experimental validation of mucin-binding in fMtkYen801

Phage binding to mucus was initially predicted *in silico* by identifying Ig-like domains in the fMtkYen801 genome using Pharokka v1.7.4 (70). The corresponding CDS was further validated with HHPred (default parameters) against the Protein Data Bank (PDB) database (71). Structural prediction of the protein was performed based on the raw amino acid sequences using ColabFold v1.5.5 combining Alphafold2 with MMseqs2 (72). The resulting PDB file was further analyzed and visualized using iCn3D (73). Protein sequences of the query and reference Ig-like domains were aligned using clustalw with default parameters, and without additional alignment trimming. Foldseek was used for structural comparisons of Ig domain topology (74).

The mucin-binding phenotype was experimentally evaluated as described previously (14,25). Namely, phage suspension was diluted to 2.5×10^3^ PFU/mL for fMtkYen801, and 2.5×10^2^ PFU/mL for fMtkYen3-01, 4mL of which was applied to LB agar plates supplemented with 1% (w/v) porcine gastric mucin, or equivalent amount of MilliQ water. Plates were incubated at RT on an orbital shaker for 30 min, followed by decanting the liquid samples, and overlaying the surfaces with o/n bacterial host mixed with 3mL LB soft agar (0.4% w/v). After overnight incubation at RT, the number of plaques, indicating the number of phage particles adhered to agar surfaces, was estimated. The experiment was performed in 12 replicates.

### The effect of mucin and multiplicity of infection (MOI) on phage-host dynamics

An overnight culture of *Y.enterocolitica* was diluted to Optical Density (OD_595_) of 0.6 (1-2 x 10^9^ CFU/mL) (Multiskan FC Microplate Reader, Thermo Scientific) and mixed with different phage concentrations to achieve MOI of 0.1, 0.01 and 0.001, respectively, in the final volume of 200 µL. The above-mentioned treatment groups were studied under 0%, 0.05%, 0.1% and 0.2% mucin supplemented conditions and incubated at 25°C, with constant agitation. The experiment was performed on Bioscreen C analyzer (Growth Curves AB Ltd, Finland), with OD_600_ measurement every 30 min for 60 h. Phage-naïve bacterial cultures, grown in LB with or without mucin supplementation, were included as positive controls. Sterile LB, supplemented with corresponding mucin concentration (0-0.2%) served as negative controls, and was used for background subtraction from wells with test conditions. At the end of experiment, 0.1x volume of chloroform was applied to phage-treated groups, settled at 4°C, and the supernatant was used for phage enumeration by drop test. All treatment groups were tested in 10 techincal replicates. One-way ANOVA followed by Tukey’s multiple comparison test was performed using GraphPad Prism version 10.0.0 for Windows, GraphPad Software (Boston, Massachusetts USA).

### The effect of mucin on phage-host dynamics under varying environmental conditions

The overnight culture of *Y. enterocolitica* was adjusted to OD_600_ of 0.6, and 10-fold diluted in 3 mL LB with or without 0.2% (w/v) mucin supplementation, resulting in final concentration of 1-2×10^8^ CFU/mL. Samples were incubated at 25°C or 37°C for 2 h. Similarly, 1×10^7^ PFU/mL phage suspension was exposed to 0% or 0.2% (w/v) mucin at varying temperatures (25°C or 37°C) for 2 h before the host infection. After 2-hour incubation, 100 µL of the bacterial samples were aliquoted in two separate Honeycomb plates, infected with 100uL of the respective phages from control and mucin treatments, and incubated for another 24 h at 25°C or 37°C, respectively. Cultures were monitored in a Bioscreen C system, under continuous agitation and optical density measurement every 30 minutes. After 24 h, phage titers and viable cell counts were evaluated. Two independent experiments were performed, with 10 replicates in each treatment group.

Similarly, phage-bacterium dynamics were tested under contrasting nutrient conditions (LB vs dH₂O) as described above, with the following modifications: bacterial cultures were infected with phage at MOI of 0.1 (1×10^7^ CFU bacteria to 1×10^6^ PFU phage) in either standard nutrient medium (LB) or sterile distilled water (dH₂O), with or without 2 h mucin pretreatment. All the incubations were performed at 25°C. At the end of the experiment, bacterial growth curves were generated based on optical density measurements performed on Bioscreen C system, following the background subtraction using cell-free medium wells with or without mucin supplementation. Phage and bacterial populations were quantified by determining plaque-forming units (PFUs) and colony-forming units (CFUs), respectively. Data was collected from 2 independent experiments, in 10 technical replicates.

### The effect of mucin on biofilm formation

Overnight bacterial suspensions were adjusted to OD_600_=0.6 and diluted 1:10 to achieve the final concentration of 1-2×10^8^ CFU/mL. 1 mL of the diluted cultures were aliquoted and harvested at 4,600 x g, 20 min, followed by resuspension of the pellets in 1X LB, 0.1X LB, dH₂O, supplemented with 0% or 0.2% (w/v) mucin. 150 µL of the above-mentioned samples (1.5 x 10^7^ CFU/well) were distributed to sterile 96-well plates, and exposed to 10 µL phage (1×10^6^ PFU/well) or LB. After static incubation at RT for 48 h, planktonic cells and residual medium were removed, plates were washed three times with sterile water and subsequently stained with 0.1% crystal violet. Following staining, plates were washed again three times with dH₂O, air-dried and de-stained with 96% ethanol. After crystal violet solubilization, 100 µL of each sample was transferred to a new 96-well plate and absorbance was measured at 595 nm (Multiskan FC Microplate Reader, Thermo Scientific). Absorbance values for each treatment group were corrected by subtracting OD_595_ of respective sterile medium with or without mucin supplementation (1X LB, 0.1X LB, dH_2_O). Each treatment was tested in 10 technical replicates from two independent experiments.

### Comparative genomics of phage resistant mutants

To generate bacteriophage-insensitive mutants (BIMs) from the ancestral *Y. enterocolitica* strain, mid-log-phase cultures (OD=0.6) were infected with phage fMtkYen801 at MOI of 0.1 (1×10^8^ CFU bacteria to 1×10^7^ PFU phage) and incubated at 25°C for 60 h under 0% or 0.2% mucin supplemented conditions. Following the incubation, serial dilutions of the samples were plated on LB agar to produce individual colonies, and incubated at RT for 48 h. Next, 10 colonies per treatment were randomly selected and purified using three successive rounds of re-streaking. Phage resistance profile of the purified colonies was confirmed using spot test, after which BIMs were stored in 15% glycerol stocks for the downstream experiments.

DNA was extracted from 1×10^9^ CFU/mL bacterial cultures using Qiagen DNeasy 96 Blood & Tissue Kit (Qiagen, Hilden, Germany). The purity and concentration of DNA was assessed on NanoDrop spectrophotometer for the single BIM isolates from 0% and 0.2% mucin treatments, as well as the ancestral strain. Next, the samples were prepared using the Oxford Nanopore Native Barcoding Kit 24 V14 (SQK-NBD114.24; Oxford Nanopore Technologies (ONT)) for library preparation and barcoding as per recommended protocol. The library was sequenced using the SQK-NBD114.24 protocol on MinKNOW v. 23.11.5, then basecalled and demultiplexed using the Dorado suite (v.0.8.3; ONT). The resulting reads were quality checked using seqkit v.0.16.0 (75) with reads < 500 bp and Quality score (Q) < 8 removed prior to further analysis. The ancestral strain was additionally commercially sequenced on DNBseq platform (BGI), with 150 paired-end protocol.

For hybrid assembly of the ancestral genome with long- and short-read data, the following pipeline was followed: Nanopore draft assembly was generated using Flye v2.9.5 (76) with quality filtered long reads. The initial assembly was polished by re-mapping raw reads back to the assembly using minimap2 (77), followed by assessment of the quality and completeness using Quast 5.2.0 (78) and Busco 5.8.2 (79), respectively. Next, Nanopore assembly was error-corrected with DNBseq short reads by alignment using BWA-MEM 0.7.18 (80), followed by polishing with Polypolish (81), and an additional round of quality control.

Annotation was performed using Prokka 1.14.6 (82). ProgressiveMauve and Blast Ring Image Generator (BRIG) (83,84) were used for the alignment of the BIM sequences to the reference strain. Plasmid marker genes and defense mechanisms were identified by PlasmidFinder 2.1 (85) and Padloc v.2.0.0 (86), respectively. Phastest (87) was used for finding the prophage regions.

Comparative genomic analysis was performed by mapping the *Y. enterocolitica* BIM sequencing reads to the ancestral genome assembly using minimap2 (77), and variant calling with Bcftools v1.21 (88). After filtering low quality variants, SnpEff v4.1k_cv3 (89) was used for variant annotation relative to the reference genome sequence. Variant Call Format (VCF) files were visualized in Integrative Genomics Viewer (IGV) (90). Gene functional and metabolic pathways were analyzed using Uniprot and KEGG (91–93), respectively.

## Acknowledgements

We would like to thank Petri Papponen for skillful help in Transmission Electron Microscopy (TEM). We are grateful to Andrew Millard for helpful discussions.

## Funding

This work was supported by Research Council of Finland (#346772, for L.-R.S and #354982 for R. P.), Centre for New Antibacterial Strategies (CANS) of the Arctic University of Norway (project ID #2520855, for G.M.F.A), and Finnish National Agency for Education (EDUFI) (TM-21-11539, for S.G.).

## Data availability

Upon publication, all sequence data will be made publicly available through GenBank, and the datasets supporting the conclusions of this article will be published in the JYX repository.

## Notes

### Competing Interest Statement

The authors have declared no competing interest.

## References

1. Linden SK, Sutton P, Karlsson NG, Korolik V, McGuckin MA. Mucins in the mucosal barrier to infection. Mucosal Immunol. 2008 May;1(3):183–97. doi:10.1038/mi.2008.5

2. Cone RA. Barrier properties of mucus. Advanced Drug Delivery Reviews. 2009 Feb 27;61(2):75–85. doi:10.1016/j.addr.2008.09.008

3. Corfield AP, Carroll D, Myerscough N, Probert CSJ. Mucins in the gastrointestinal tract in health and disease. FBL. 2001 Oct 1;6(3):3. doi:10.2741/corfield

4. Ohneck EJ, Arivett BA, Fiester SE, Wood CR, Metz ML, Simeone GM, et al. Mucin acts as a nutrient source and a signal for the differential expression of genes coding for cellular processes and virulence factors in Acinetobacter baumannii. PLOS ONE. 2018 Jan 8;13(1):e0190599. doi:10.1371/journal.pone.0190599

5. Schroeder BO. Fight them or feed them: how the intestinal mucus layer manages the gut microbiota. Gastroenterology Report. 2019 Feb 1;7(1):3–12. doi:10.1093/gastro/goy052

6. Sicard JF, Le Bihan G, Vogeleer P, Jacques M, Harel J. Interactions of Intestinal Bacteria with Components of the Intestinal Mucus. Frontiers in Cellular and Infection Microbiology [Internet]. 2017 [cited 2023 Apr 18];7. Available from: https://www.frontiersin.org/articles/10.3389/fcimb.2017.00387

7. Brockhausen I, Falconer D, Sara S. Relationships between bacteria and the mucus layer. Carbohydrate Research. 2024 Dec 1;546:109309. doi:10.1016/j.carres.2024.109309

8. Bakshani CR, Morales-Garcia AL, Althaus M, Wilcox MD, Pearson JP, Bythell JC, et al. Evolutionary conservation of the antimicrobial function of mucus: a first defence against infection. npj Biofilms Microbiomes. 2018 Jul 4;4(1):14. doi:10.1038/s41522-018-0057-2

9. Zhou X, Wu Y, Zhu Z, Lu C, Zhang C, Zeng L, et al. Mucosal immune response in biology, disease prevention and treatment. Sig Transduct Target Ther. 2025 Jan 8;10(1):7. doi:10.1038/s41392-024-02043-4

10. Hansson GC. Mucins and the Microbiome. Annual Review of Biochemistry. 2020 Jun 20;89(Volume 89, 2020):769–93. doi:10.1146/annurev-biochem-011520-105053

11. Fang J, Wang H, Zhou Y, Zhang H, Zhou H, Zhang X. Slimy partners: the mucus barrier and gut microbiome in ulcerative colitis. Exp Mol Med. 2021 May;53(5):772–87. doi:10.1038/s12276-021-00617-8

12. Rothschild-Rodriguez D, Hedges M, Kaplan M, Karav S, Nobrega FL. Phage-encoded carbohydrate-interacting proteins in the human gut. Front Microbiol. 2023 Jan 6;13. doi:10.3389/fmicb.2022.1083208

13. Fraser JS, Yu Z, Maxwell KL, Davidson AR. Ig-Like Domains on Bacteriophages: A Tale of Promiscuity and Deceit. Journal of Molecular Biology. 2006 Jun 2;359(2):496–507. doi:10.1016/j.jmb.2006.03.043

14. Barr JJ, Auro R, Furlan M, Whiteson KL, Erb ML, Pogliano J, et al. Bacteriophage adhering to mucus provide a non–host-derived immunity. Proceedings of the National Academy of Sciences. 2013 Jun 25;110(26):10771–6. doi:10.1073/pnas.1305923110

15. Barr JJ, Auro R, Sam-Soon N, Kassegne S, Peters G, Bonilla N, et al. Subdiffusive motion of bacteriophage in mucosal surfaces increases the frequency of bacterial encounters. Proceedings of the National Academy of Sciences. 2015 Nov 3;112(44):13675–80. doi:10.1073/pnas.1508355112

16. Joiner KL, Baljon A, Barr J, Rohwer F, Luque A. Impact of bacteria motility in the encounter rates with bacteriophage in mucus. Sci Rep. 2019 Nov 11;9(1):16427. doi:10.1038/s41598-019-52794-2

17. Chin WH, Kett C, Cooper O, Müseler D, Zhang Y, Bamert RS, et al. Bacteriophages evolve enhanced persistence to a mucosal surface. Proceedings of the National Academy of Sciences. 2022 Jul 5;119(27):e2116197119. doi:10.1073/pnas.2116197119

18. Sundberg LR, Rantanen N, De Freitas Almeida GM. Mucosal Environment Induces Phage Susceptibility in *Streptococcus mutans*. PHAGE. 2022 Sep 1;3(3):128–35. doi:10.1089/phage.2022.0021

19. Pacios O, Blasco L, Ortiz Cartagena C, Bleriot I, Fernández-García L, López M, et al. Molecular studies of phages-Klebsiella pneumoniae in mucoid environment: innovative use of mucolytic agents prior to the administration of lytic phages. Front Microbiol. 2023 Oct 11;14. doi:10.3389/fmicb.2023.1286046

20. Núñez-Sánchez MA, Colom J, Walsh L, Buttimer C, Bolocan AS, Pang R, et al. Characterizing Phage-Host Interactions in a Simplified Human Intestinal Barrier Model. Microorganisms. 2020 Sep;8(9):9. doi:10.3390/microorganisms8091374

21. Green SI, Gu Liu C, Yu X, Gibson S, Salmen W, Rajan A, et al. Targeting of Mammalian Glycans Enhances Phage Predation in the Gastrointestinal Tract. mBio. 2021 Feb 9;12(1):10.1128/mbio.03474-20. doi:10.1128/mbio.03474-20

22. Jakin Lazar J, Šimunović K, Dogša I, Mandić Mulec I, Middelboe M, Dragoš A. Distinct effects of mucin on phage-host interactions in model systems of beneficial and pathogenic bacteria. Arch Virol. 2025 May 20;170(6):133. doi:10.1007/s00705-025-06322-5

23. Almeida GM de F, Ravantti J, Grdzelishvili N, Kakabadze E, Bakuradze N, Javakhishvili E, et al. Relevance of the bacteriophage adherence to mucus model for Pseudomonas aeruginosa phages. Microbiology Spectrum. 2024 Jun 24;12(8):e03520–23. doi:10.1128/spectrum.03520-23

24. de Freitas Almeida GM, Hoikkala V, Ravantti J, Rantanen N, Sundberg LR. Mucin induces CRISPR-Cas defense in an opportunistic pathogen. Nat Commun. 2022 Jun 25;13(1):3653. doi:10.1038/s41467-022-31330-3

25. Almeida GMF, Laanto E, Ashrafi R, Sundberg LR. Bacteriophage Adherence to Mucus Mediates Preventive Protection against Pathogenic Bacteria. Martiny JBH, editor. mBio. 2019 Dec 24;10(6):e01984–19. doi:10.1128/mBio.01984-19

26. Coelho LFL, de Souza Terceti M, Neto SPL, Amaral RP, dos Santos ALC, Gozzi WP, et al. Mucosal-adapted bacteriophages as a preventive strategy for a lethal Pseudomonas aeruginosa challenge in mice. Commun Biol. 2025 Jan 6;8(1):1–10. doi:10.1038/s42003-024-07269-0

27. Yen M, Cairns LS, Camilli A. A cocktail of three virulent bacteriophages prevents Vibrio cholerae infection in animal models. Nat Commun. 2017 Feb 1;8(1):14187. doi:10.1038/ncomms14187

28. Wu J, Fu K, Hou C, Wang Y, Ji C, Xue F, et al. Bacteriophage defends murine gut from Escherichia coli invasion via mucosal adherence. Nat Commun. 2024 Jun 4;15(1):4764. doi:10.1038/s41467-024-48560-2

29. Fu K, Cui J, Li Y, Zhang Y, Wang Y, Wu J, et al. Escherichia coli phage ΦPNJ-9 adheres to mucus via a variant Hoc protein. J Virol. 99(2):e01789–24. doi:10.1128/jvi.01789-24 PubMed PMID: 39723818; PubMed Central PMCID: PMC11853027.

30. Xue Y, Zhai S, Wang Z, Ji Y, Wang G, Wang T, et al. The Yersinia Phage X1 Administered Orally Efficiently Protects a Murine Chronic Enteritis Model Against Yersinia enterocolitica Infection. Front Microbiol. 2020;11:351. doi:10.3389/fmicb.2020.00351 PubMed PMID: 32210942; PubMed Central PMCID: PMC7067902.

31. Murray CJL, Ikuta KS, Sharara F, Swetschinski L, Aguilar GR, Gray A, et al. Global burden of bacterial antimicrobial resistance in 2019: a systematic analysis. The Lancet. 2022 Feb 12;399(10325):629–55. doi:10.1016/S0140-6736(21)02724-0 PubMed PMID: 35065702.

32. Grygiel-Górniak B. Current Challenges in Yersinia Diagnosis and Treatment. Microorganisms. 2025 May 15;13(5):1133. doi:10.3390/microorganisms13051133 PubMed PMID: 40431305; PubMed Central PMCID: PMC12114158.

33. EU summary report on zoonoses, zoonotic agents and food-borne outbreaks 2013 | EFSA [Internet]. 2015 [cited 2025 Jul 10]. Available from: https://www.efsa.europa.eu/en/efsajournal/pub/3991

34. Li C, Gölz G, Alter T, Barac A, Hertwig S, Riedel C. Prevalence and Antimicrobial Resistance of Yersinia enterocolitica in Retail Seafood. J Food Prot. 2018 Mar 1;81(3):497–501. doi:10.4315/0362-028X.JFP-17-357 PubMed PMID: 29474145.

35. Bonardi S, Paris A, Bassi L, Salmi F, Bacci C, Riboldi E, et al. Detection, semiquantitative enumeration, and antimicrobial susceptibility of Yersinia enterocolitica in pork and chicken meats in Italy. J Food Prot. 2010 Oct;73(10):1785–92. doi:10.4315/0362-028x-73.10.1785 PubMed PMID: 21067665.

36. Goladze S, Patpatia S, Tuomala H, Ylänne M, Gachechiladze N, de Oliveira Patricio D, et al. Isolation and characterization of Yersinia phage fMtkYen3-01. Arch Virol. 2024 Oct 19;169(11):226. doi:10.1007/s00705-024-06149-6

37. Skurnik M, Bengoechea JA. The biosynthesis and biological role of lipopolysaccharide O-antigens of pathogenic *Yersiniae*. Carbohydrate Research. 2003 Nov 14;Bacterial Antigens and Vaccines338(23):2521–9. doi:10.1016/S0008-6215(03)00305-7

38. Bengoechea JA, Zhang L, Toivanen P, Skurnik M. Regulatory network of lipopolysaccharide O-antigen biosynthesis in Yersinia enterocolitica includes cell envelope-dependent signals. Molecular Microbiology. 2002;44(4):1045–62. doi:10.1046/j.1365-2958.2002.02940.x

39. Leon-Velarde CG, Happonen L, Pajunen M, Leskinen K, Kropinski AM, Mattinen L, et al. Yersinia enterocolitica-Specific Infection by Bacteriophages TG1 and ϕR1-RT Is Dependent on Temperature-Regulated Expression of the Phage Host Receptor OmpF. Applied and Environmental Microbiology. 2016 Sep;82(17):5340–53. doi:10.1128/AEM.01594-16

40. Sauvage E, Kerff F, Terrak M, Ayala JA, Charlier P. The penicillin-binding proteins: structure and role in peptidoglycan biosynthesis. FEMS Microbiology Reviews. 2008 Mar 1;32(2):234–58. doi:10.1111/j.1574-6976.2008.00105.x

41. Srivastava D, Waters CM. A Tangled Web: Regulatory Connections between Quorum Sensing and Cyclic Di-GMP. J Bacteriol. 2012 Sep;194(17):4485–93. doi:10.1128/JB.00379-12 PubMed PMID: 22661686; PubMed Central PMCID: PMC3415487.

42. Wang M, Wang J, Li T, Bao X, Li P, Zhang X, et al. Penicillin-binding protein 1b encoded by mrcB gene mediates the enhancement of biofilm formation by subinhibitory concentrations of cefotaxime in monophasic Salmonella Typhimurium strain SH16SP46. FEMS Microbiology Letters. 2023 Jan 1;370:fnad021. doi:10.1093/femsle/fnad021

43. Jenal U, Reinders A, Lori C. Cyclic di-GMP: second messenger extraordinaire. Nat Rev Microbiol. 2017 May;15(5):271–84. doi:10.1038/nrmicro.2016.190

44. Akhtar AA, Turner DPJ. The role of bacterial ATP-binding cassette (ABC) transporters in pathogenesis and virulence: Therapeutic and vaccine potential. Microbial Pathogenesis. 2022 Oct 1;171:105734. doi:10.1016/j.micpath.2022.105734

45. Skurnik M, Biedzka-Sarek M, Lübeck PS, Blom T, Bengoechea JA, Pérez-Gutiérrez C, et al. Characterization and Biological Role of the O-Polysaccharide Gene Cluster of Yersinia enterocolitica Serotype O:9. J Bacteriol. 2007 Oct;189(20):7244–53. doi:10.1128/JB.00605-07 PubMed PMID: 17693522; PubMed Central PMCID: PMC2168460.

46. Skurnik M, Peippo A, Ervelä E. Characterization of the O-antigen gene clusters of Yersinia pseudotuberculosis and the cryptic O-antigen gene cluster of Yersinia pestis shows that the plague bacillus is most closely related to and has evolved from Y. pseudotuberculosis serotype O:1b. Molecular Microbiology. 2000;37(2):316–30. doi:10.1046/j.1365-2958.2000.01993.x

47. Zhang L, Radziejewska-Lebrecht J, Krajewska-Pietrasik D, Toivanen P, Skurnik M. Molecular and chemical characterization of the lipopolysaccharide O-antigen and its role in the virulence of Yersinia enterocolitica serotype O:8. Molecular Microbiology. 1997;23(1):63–76. doi:10.1046/j.1365-2958.1997.1871558.x

48. Zhang L, Toivanen P, Skurnik M. The gene cluster directing O-antigen biosynthesis in Yersinia enterocolitica serotype O:8: identification of the genes for mannose and galactose biosynthesis and the gene for the O-antigen polymerase. Microbiology. 1996;142(2):277–88. doi:10.1099/13500872-142-2-277

49. Carroll-Portillo A, Lin HC. Exploring Mucin as Adjunct to Phage Therapy. Microorganisms. 2021 Feb 28;9(3):509. doi:10.3390/microorganisms9030509 PubMed PMID: 33670927; PubMed Central PMCID: PMC7997181.

50. Sathaliyawala T, Islam MZ, Li Q, Fokine A, Rossmann MG, Rao VB. Functional Analysis of the Highly Antigenic Outer Capsid Protein, Hoc, a Virus Decoration Protein from T4-like Bacteriophages. Mol Microbiol. 2010 Jul;77(2):444–55. doi:10.1111/j.1365-2958.2010.07219.x PubMed PMID: 20497329; PubMed Central PMCID: PMC2909354.

51. Fraser JS, Yu Z, Maxwell KL, Davidson AR. Ig-like domains on bacteriophages: a tale of promiscuity and deceit. J Mol Biol. 2006 Jun 2;359(2):496–507. doi:10.1016/j.jmb.2006.03.043 PubMed PMID: 16631788.

52. Carroll-Portillo A, Rumsey KN, Braun CA, Lin DM, Coffman CN, Alcock JA, et al. Mucin and Agitation Shape Predation of Escherichia coli by Lytic Coliphage. Microorganisms. 2023 Feb;11(2):2. doi:10.3390/microorganisms11020508

53. Wu J, Fu K, Hou C, Wang Y, Ji C, Xue F, et al. Bacteriophage defends murine gut from Escherichia coli invasion via mucosal adherence. Nat Commun. 2024 Jun 4;15(1):4764. doi:10.1038/s41467-024-48560-2

54. Shan J, Ramachandran A, Thanki AM, Vukusic FBI, Barylski J, Clokie MRJ. Bacteriophages are more virulent to bacteria with human cells than they are in bacterial culture; insights from HT-29 cells. Sci Rep. 2018 Mar 23;8(1):5091. doi:10.1038/s41598-018-23418-y

55. Bath J, Bjånes E, Goekeri C, Hsiao J, Uzun D, Nouailles G, et al. Mucins protect against Streptococcus pneumoniae virulence by suppressing pneumolysin expression. J Clin Invest. 134(19):e182769. doi:10.1172/JCI182769 PubMed PMID: 39172520; PubMed Central PMCID: PMC11444151.

56. Wheeler KM, Cárcamo-Oyarce G, Turner BS, Dellos-Nolan S, Co JY, Lehoux S, et al. Mucin glycans attenuate the virulence of Pseudomonas aeruginosa in infection. Nat Microbiol. 2019 Dec 1;4(12):2146–54. doi:10.1038/s41564-019-0581-8 PubMed PMID: 31611643; PubMed Central PMCID: PMC7157942.

57. Martens EC, Chiang HC, Gordon JI. Mucosal Glycan Foraging Enhances Fitness and Transmission of a Saccharolytic Human Gut Bacterial Symbiont. Cell Host & Microbe. 2008 Nov 13;4(5):447–57. doi:10.1016/j.chom.2008.09.007 PubMed PMID: 18996345.

58. Tailford LE, Crost EH, Kavanaugh D, Juge N. Mucin glycan foraging in the human gut microbiome. Front Genet. 2015 Mar 19;6. doi:10.3389/fgene.2015.00081

59. Etzold S, Juge N. Structural insights into bacterial recognition of intestinal mucins. Current Opinion in Structural Biology. 2014 Oct 1;28:23–31. doi:10.1016/j.sbi.2014.07.002

60. Mantle M, Rombough C. Growth in and breakdown of purified rabbit small intestinal mucin by Yersinia enterocolitica. Infect Immun. 1993 Oct;61(10):4131–8. doi:10.1128/iai.61.10.4131-4138.1993 PubMed PMID: 8406802; PubMed Central PMCID: PMC281135.

61. Fara E, Raach B, Cavallaro A, Fink J, Ramesh D, Guo Y, et al. Phage-mediated lysis increases growth rate of surviving bacterial cells.

62. Latka A, Maciejewska B, Majkowska-Skrobek G, Briers Y, Drulis-Kawa Z. Bacteriophage-encoded virion-associated enzymes to overcome the carbohydrate barriers during the infection process. Appl Microbiol Biotechnol. 2017 Apr 1;101(8):3103–19. doi:10.1007/s00253-017-8224-6

63. Clark A, Mach N. The gut mucin-microbiota interactions: a missing key to optimizing endurance performance. Front Physiol. 2023 Nov 22;14:1284423. doi:10.3389/fphys.2023.1284423 PubMed PMID: 38074323; PubMed Central PMCID: PMC10703311.

64. Arias SL, van Wijngaarden EW, Balint D, Jones J, Crawford CC, Shukla PJ, et al. Environmental factors drive bacterial degradation of gastrointestinal mucus. NPJ Biofilms Microbiomes. 2025 Jul 16;11:133. doi:10.1038/s41522-025-00741-7 PubMed PMID: 40670389; PubMed Central PMCID: PMC12267731.

65. Elzinga J, Narimatsu Y, de Haan N, Clausen H, de Vos WM, Tytgat HLP. Binding of Akkermansia muciniphila to mucin is O-glycan specific. Nat Commun. 2024 May 29;15(1):4582. doi:10.1038/s41467-024-48770-8

66. Zhang L, Toivanen P, Skurnik M. The gene cluster directing O-antigen biosynthesis in Yersinia enterocolitica serotype O:8: identification of the genes for mannose and galactose biosynthesis and the gene for the O-antigen polymerase. Microbiology. 1996;142(2):277–88. doi:10.1099/13500872-142-2-277

67. Pajunen M, Kiljunen S, Skurnik M. Bacteriophage φYeO3-12, Specific for Yersinia enterocolitica Serotype O:3, Is Related to Coliphages T3 and T7. J Bacteriol. 2000 Sep;182(18):5114–20. doi:10.1128/jb.182.18.5114-5120.2000 PubMed PMID: 10960095; PubMed Central PMCID: PMC94659.

68. Salem M, Skurnik M. Genomic Characterization of Sixteen Yersinia enterocolitica-Infecting Podoviruses of Pig Origin. Viruses. 2018 Apr;10(4):4. doi:10.3390/v10040174

69. Bengoechea JA, Najdenski H, Skurnik M. Lipopolysaccharide O antigen status of *Yersinia enterocolitica* O:8 is essential for virulence and absence of O antigen affects the expression of other *Yersinia* virulence factors. Molecular Microbiology. 2004 Apr;52(2):451–69. doi:10.1111/j.1365-2958.2004.03987.x

70. Bouras G, Nepal R, Houtak G, Psaltis AJ, Wormald PJ, Vreugde S. Pharokka: a fast scalable bacteriophage annotation tool. Bioinformatics. 2023 Jan 1;39(1):btac776. doi:10.1093/bioinformatics/btac776

71. Zimmermann L, Stephens A, Nam SZ, Rau D, Kübler J, Lozajic M, et al. A Completely Reimplemented MPI Bioinformatics Toolkit with a New HHpred Server at its Core. Journal of Molecular Biology. 2018 Jul 20;Computation Resources for Molecular Biology430(15):2237–43. doi:10.1016/j.jmb.2017.12.007

72. Mirdita M, Schütze K, Moriwaki Y, Heo L, Ovchinnikov S, Steinegger M. ColabFold: making protein folding accessible to all. Nat Methods. 2022 Jun;19(6):679–82. doi:10.1038/s41592-022-01488-1

73. Wang J, Youkharibache P, Zhang D, Lanczycki CJ, Geer RC, Madej T, et al. iCn3D, a web-based 3D viewer for sharing 1D/2D/3D representations of biomolecular structures. Bioinformatics. 2020 Jan 1;36(1):131–5. doi:10.1093/bioinformatics/btz502

74. van Kempen M, Kim SS, Tumescheit C, Mirdita M, Lee J, Gilchrist CLM, et al. Fast and accurate protein structure search with Foldseek. Nat Biotechnol. 2024 Feb;42(2):243–6. doi:10.1038/s41587-023-01773-0

75. Shen W, Le S, Li Y, Hu F. SeqKit: A Cross-Platform and Ultrafast Toolkit for FASTA/Q File Manipulation. PLOS ONE. 2016 Oct 5;11(10):e0163962. doi:10.1371/journal.pone.0163962

76. Kolmogorov M, Yuan J, Lin Y, Pevzner PA. Assembly of long, error-prone reads using repeat graphs. Nat Biotechnol. 2019 May;37(5):540–6. doi:10.1038/s41587-019-0072-8

77. Li H. Minimap2: pairwise alignment for nucleotide sequences. Bioinformatics. 2018 Sep 15;34(18):3094–100. doi:10.1093/bioinformatics/bty191

78. Gurevich A, Saveliev V, Vyahhi N, Tesler G. QUAST: quality assessment tool for genome assemblies. Bioinformatics. 2013 Apr 15;29(8):1072–5. doi:10.1093/bioinformatics/btt086

79. Simão FA, Waterhouse RM, Ioannidis P, Kriventseva EV, Zdobnov EM. BUSCO: assessing genome assembly and annotation completeness with single-copy orthologs. Bioinformatics. 2015 Oct 1;31(19):3210–2. doi:10.1093/bioinformatics/btv351

80. Li H, Durbin R. Fast and accurate short read alignment with Burrows–Wheeler transform. Bioinformatics. 2009 Jul 15;25(14):1754–60. doi:10.1093/bioinformatics/btp324

81. Wick RR, Holt KE. Polypolish: Short-read polishing of long-read bacterial genome assemblies. PLOS Computational Biology. 2022 Jan 24;18(1):e1009802. doi:10.1371/journal.pcbi.1009802

82. Seemann T. Prokka: rapid prokaryotic genome annotation. Bioinformatics. 2014 Jul 15;30(14):2068–9. doi:10.1093/bioinformatics/btu153

83. progressiveMauve: Multiple Genome Alignment with Gene Gain, Loss and Rearrangement | PLOS One [Internet]. [cited 2025 Jul 11]. Available from: https://journals.plos.org/plosone/article?id=10.1371/journal.pone.0011147

84. Alikhan NF, Petty NK, Ben Zakour NL, Beatson SA. BLAST Ring Image Generator (BRIG): simple prokaryote genome comparisons. BMC Genomics. 2011 Aug 8;12(1):402. doi:10.1186/1471-2164-12-402

85. Carattoli A, Zankari E, García-Fernández A, Voldby Larsen M, Lund O, Villa L, et al. In silico detection and typing of plasmids using PlasmidFinder and plasmid multilocus sequence typing. Antimicrob Agents Chemother. 2014 Jul;58(7):3895–903. doi:10.1128/AAC.02412-14 PubMed PMID: 24777092; PubMed Central PMCID: PMC4068535.

86. Payne LJ, Meaden S, Mestre MR, Palmer C, Toro N, Fineran PC, et al. PADLOC: a web server for the identification of antiviral defence systems in microbial genomes. Nucleic Acids Research. 2022 Jul 5;50(W1):W541–50. doi:10.1093/nar/gkac400

87. Wishart DS, Han S, Saha S, Oler E, Peters H, Grant JR, et al. PHASTEST: faster than PHASTER, better than PHAST. Nucleic Acids Research. 2023 Jul 5;51(W1):W443–50. doi:10.1093/nar/gkad382

88. Danecek P, Bonfield JK, Liddle J, Marshall J, Ohan V, Pollard MO, et al. Twelve years of SAMtools and BCFtools. Gigascience. 2021 Feb 16;10(2):giab008. doi:10.1093/gigascience/giab008 PubMed PMID: 33590861; PubMed Central PMCID: PMC7931819.

89. Cingolani P, Platts A, Wang LL, Coon M, Nguyen T, Wang L, et al. A program for annotating and predicting the effects of single nucleotide polymorphisms, SnpEff: SNPs in the genome of Drosophila melanogaster strain w^1118^; iso-2; iso-3. Fly. 2012 Apr;6(2):80–92. doi:10.4161/fly.19695

90. Robinson JT, Thorvaldsdottir H, Turner D, Mesirov JP. igv.js: an embeddable JavaScript implementation of the Integrative Genomics Viewer (IGV). Bioinformatics. 2023 Jan 1;39(1):btac830. doi:10.1093/bioinformatics/btac830

91. The UniProt Consortium. UniProt: the Universal Protein Knowledgebase in 2023. Nucleic Acids Research. 2023 Jan 6;51(D1):D523–31. doi:10.1093/nar/gkac1052

92. Kanehisa M, Furumichi M, Sato Y, Ishiguro-Watanabe M, Tanabe M. KEGG: integrating viruses and cellular organisms. Nucleic Acids Research. 2021 Jan 8;49(D1):D545–51. doi:10.1093/nar/gkaa970

93. Kanehisa M, Goto S. KEGG: Kyoto Encyclopedia of Genes and Genomes. Nucleic Acids Research. 2000 Jan 1;28(1):27–30. doi:10.1093/nar/28.1.27

