## Supporting Information 1 for "Mucin modulates phage infection dynamics and biofilm formation in enteropathogenic *Yersinia enterocolitica*"

**Phenotypic and genomic characterization of *Yersinia* phage fMtkYen801**

**Materials and Methods**

**Phage hunting and propagation**

Phage fMtkYen801 was isolated from water samples collected from Mtkvari River, Georgia. Briefly, 1 mL filtered water sample was added to 9 mL LB and 0.1 mL log phase enrichment host, and incubated at RT, 120 RPM for 24 h. Bacterial cells were then removed by centrifugation (4500 RPM, 10 min) and filtered through 0.22 µm membrane (Ministart®, Sartorius, Göttingen, Germany). Following three rounds of plaque purification, phages were propagated using the Gabrichevsky method (1), supplemented with 8% (w/v) sucrose and stored at 4°C until further use.

**Transmission electron microscopy (TEM)**

To study phage fMtkYen801 structure using transmission electron microscopy (TEM), PEG-precipitated sample was purified by glycerol gradient ultracentrifugation, harvested at 20,000 rpm, 4°C, 2 h and dissolved in 50 uL 0.1 M ammonium acetate, as previously described (2). Next, phage particles were sedimented on carbon coated copper grids, incubated for 30 sec, and negatively stained with 2% uranyl acetate. Specimens were studied using JEOL JEM-1400 transmission electron microscope (JEOL Ltd., Tokyo, Japan) under 80 kV. Measurement data was collected from ten virions, and mean values and standard deviations were calculated.

**DNA extraction, sequencing, and analysis**

Phage DNAs were extracted from 2 × 10^10^ PFU/mL stock using Maxwell® RSC Viral TNA puriﬁcation kit (Cat no AS1330, Promega Corporation, Madison, WI, USA), according to the manufacturer’s instructions. DNA samples were sequenced on Illumina HiSeq platform at Novogene, with 150 bp paired-end protocol (Cambridge, UK). Phage genome was assembled using A5 pipeline (Version A5-miseq 20160825) (3), with 250,000 raw reads, resulting in 35x sequencing coverage. The assembly was veriﬁed by mapping the original reads back to the genome with Geneious Prime 2023.0.4 assembler (Biomatters Ltd., Auckland, New Zealand). DNA termini and phage packaging mechanisms were evaluated using PhageTerm (4). Genome annotation was performed with Pharokka (v1.7.3) (5). Taxonomy classification was performed using Taxmyphage (6). Pairwise intergenomic distances between fMtkYen801 and other closely related phages available in GenBank database were calculated by VIRIDIC (GeneBank accessions: MK733259.1, MK733260.1, MK733261.1, OR620963.1, OL631484.1, PQ352032.1) (7). Viral proteomic tree was constructed using ViPTree server (8). Evolutionary distances between fMtkYen801 and related *Caudoviricetes* phages were calculated using VipTreeGen v1.1.3 based on pairwise tBLASTx comparisons and the resulting phylogenetic tree was visualized using Interactive Tree of Life (iToL) (9).

**Time kill assay**

The *in vitro* infectivity of fMtkYen801 was assessed by time-kill assay as previously described (10–12). The experiment covered 42 Yersinia strains, representing 4 *Yersinia* species and 18 serotypes (Supporting table 1), and was performed in triplicates. Ten µL of phage lysate containing 10^8^ Plaque Forming Units PFU/mL (10^6^ PFU/well) was added to wells of a honeycomb plate containing 200 µL of 1:20 diluted overnight (o/n) host culture, resulting in about 1x10^8^ CFU inoculum, and phage-to-bacteria ratio (MOI) of 0.01**.** Bacteria without phage infection were used as positive controls, while sterile LB served as a negative control. Plates underwent incubation at 25°C, with continuous shaking for 10 h, and optical density (OD_600_) of planktonic bacterial cultures was measured every 30 minutes using Bioscreen C analyzer (Growth Curves AB Ltd, Finland). For each time point, the mean optical density and standard deviation (SD) were calculated. Phage infection dynamics in the tested bacteria were evaluated by calculating percent reduction of bacterial growth throughout the experiment, relative to their positive controls. Phage infection efficiency was determined based on OD_600_ measurements: phage-exposed samples with OD_600_ reduction with ≥50% relative to the bacterial control were classified as infectious; samples with 30-50% reduction were deemed partially infectious, and those with less than 30% were considered noninfectious (12). Data analysis was done in GraphPad Prism 10.1.0 (GraphPad Software Inc.; San Diego, CA, USA).

**Efficiency of plating and the role of bacterial LPS for phage infectivity**

To evaluate the ability of phage fMtkYen801 to adsorb to and replicate in engineered LPS mutant *Y. enterocolitica* O:8 strains (YeO8c-WbcEGB, 8081-c-R2) (13), and to determine the Efficiency of Plating (EOP) on the selected *Yersinia* strains (*Y. enterocolitica* 6737/80, 3604/80, CDCA2635, TAMU, wa+, 15712/83, 17869/83; Supporting Table 1), spot test was used. Briefly, 10 µL drops of 10^0^-10^6^ PFU/mL phage lysates were pipetted on the respective bacterial lawn, and produced phage plaques were enumerated after incubation at 25°C for 24 h. EOP was calculated as the ratio of average PFU on each target bacteria to PFU on the isolation host (14). Both experiments were carried out in triplicates.

**pH and thermal stability**

To test phage stability in different environmental conditions, its viability was monitored under various pH and temperature ranges. Briefly, phage lysate with an initial titer of 1x10^6^ PFU/mL was exposed to LB with pH levels from 2.0 to 12.0, incubated at 25°C, and sampled after 3 h. Similarly, viable phage particles were estimated after 3 h incubation at 25, 37, 40 or 65°C. To analyze phage stability after treatments, phage titers were determined by drop tests. PFU numbers at pH 7.0 and 25°C conditions were used as controls. All assays were performed in triplicates.

**Results**

**Morphology, Host Range, and Biological Properties**

Following the ultracentrifugation and PEG precipitation, phage fMtkYen801 morphology was analyzed by transmission electron microscopy, which revealed that fMtkYen801 possesses 113 nm ± 4 nm (n = 10) long contractile tail and a capsid of 81 nm ± 40 nm (n = 10) in diameter (Fig. S1A), classifying the phage as myovirus within the class *Caudoviricetes*.

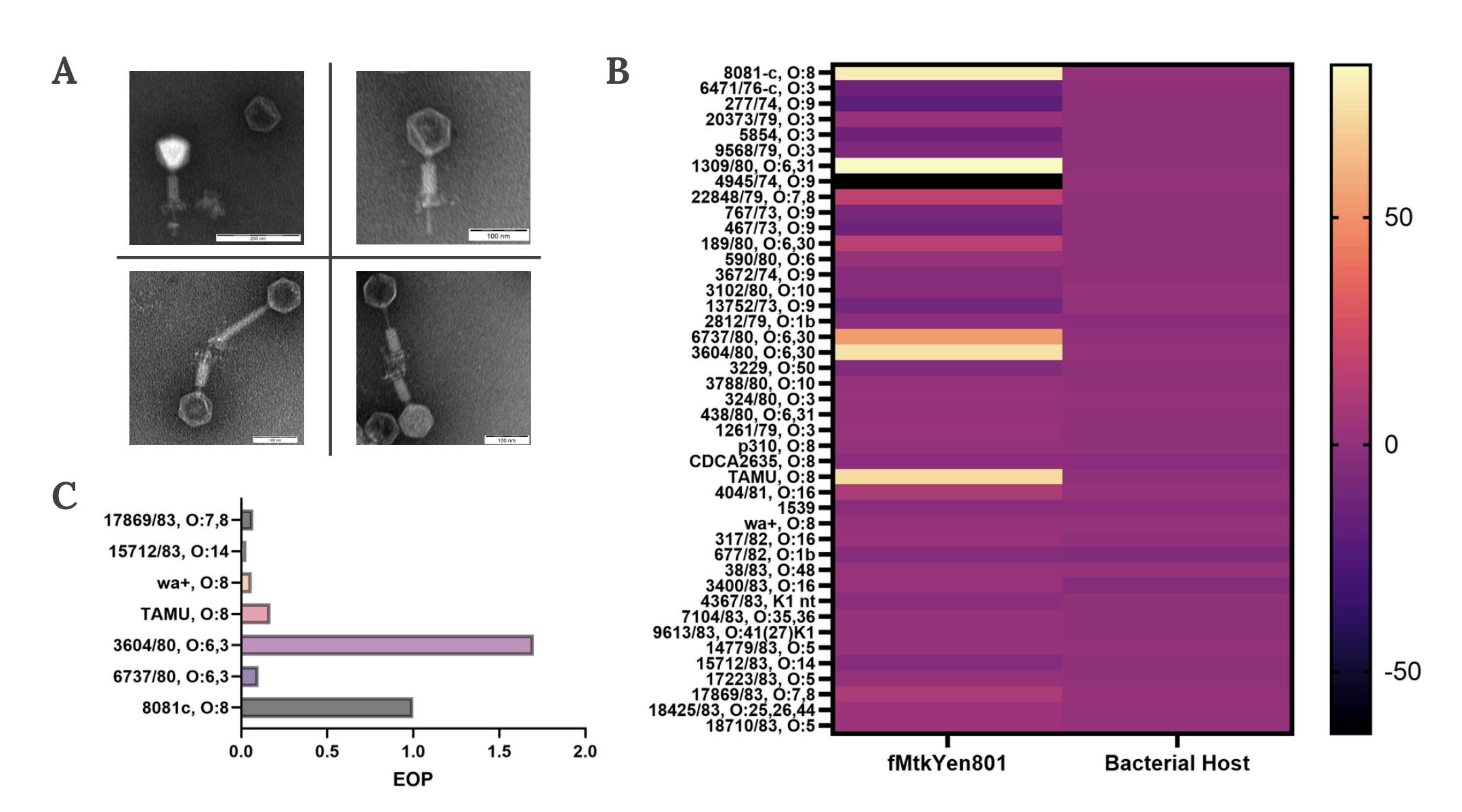

Fig. S1. Morphology, host range, and Efficiency of Plating (EOP) of *Yersinia* Phage fMtkYen801. (A) Transmission Electron Microscopy (TEM) image of fMtkYen801, showing the contractile 113 nm long tail and a capsid of 81 nm in diameter (B) Host range assessment against 42 *Yersinia* strains, showing lytic activity of fMtkYen801 against 9.3% of the tested samples. The light color indicates the sensitivity of the bacteria (C) EOP on the susceptible *Y. enterocolitica* strains

To detect the phage host specificity, a set of 42 *Yersinia* strains, including *Y.* *enterocolitica* and *Y.* *pseudotuberculosis* were challenged with fMtkYen801 in liquid medium, and optical density was monitored for 10 h, at 30 min intervals. The end-point measurements revealed that 4 (9.5 %) of the tested strains (serotypes O:8, O:6 and O:6,30) were susceptible to phage infection (Fig. S1B), with the highest Efficiency of Plating (EOP=1.7) on *Y. enterocolitica* 3605/80 (O:6,30), exceeding the EOP of the isolation host (EOP=1.0) (Fig. S1C).

To identify whether the phage used the LPS O-antigen as a receptor, two LPS mutant strains of *Y. enterocolitica* serotype O:8 (15) were evaluated using spot test, with the original host (8081-c) and its ancestral strain (8081) as references. The assay confirmed the preliminary hypothesis, as both the rough (YeO8-c-R2) and semi-rough (YeO8c-WbcEGB) mutants showed full resistance to phage fMtkYen801 infection (Fig. S2). While the rough strain is missing the complete O-polysaccharide, the semi-rough has a single O-unit on the LPS. Thus, the phage requires a polymer of O-units as a receptor.

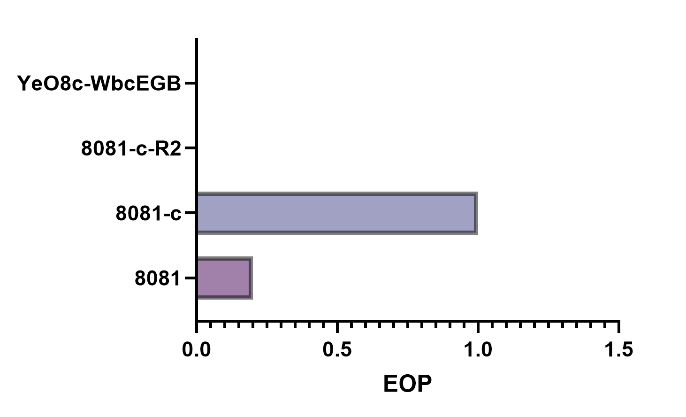

Fig. S2. Efficiency of plating (EOP) on the original host (8081c), its ancestral strain (8081) and LPS mutants (8081-c-R2, YenO8c-WbcEGB), showing resistance on O-antigen deficient variants

To estimate phage viability in various environmental conditions, pH and thermal stability of fMtkYen801 was tested. The phage could not tolerate highly acidic conditions and temperatures above 37°C, however remained stable between pH 5.0 and 9.0 and at temperatures from 25°C to 37°C for 3 hours (Fig. S3). The findings indicate that while phage fMtkYen801 remains active at physiological temperature, encapsulation or coating strategies may be necessary to protect phage particles from acidic gastric environment (16).

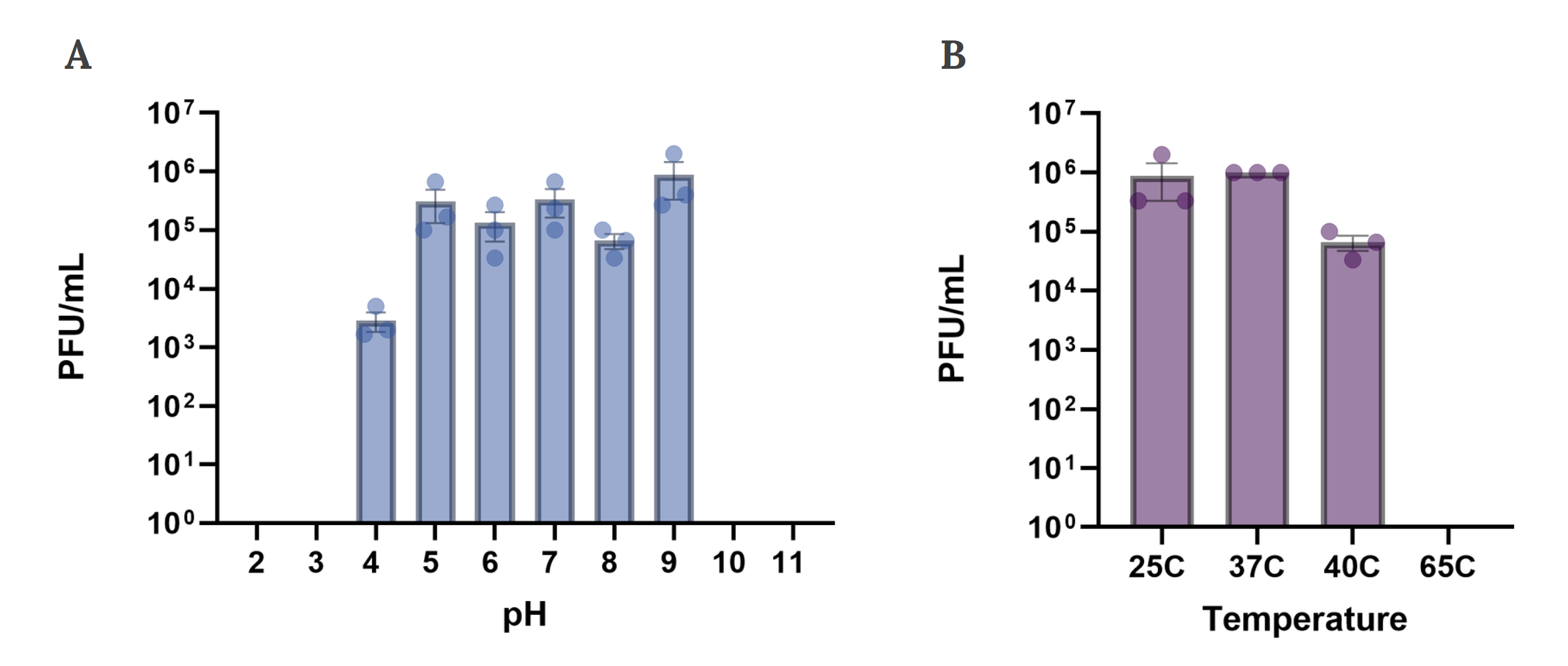

Fig. S3. Phage fMtkYen801 stability under varying pH and temperature conditions. (A) pH stability assay showing sustained phage titers at pH 5-9 (B) Temperature stability assay indicating the highest phage stability between 25°C and 37°C.

**Genomic Features and Comparative Analysis**

Pharokka (v1.7.3) identified 130 coding sequences in the *Yersinia* phage fMtkYen801 genome, with functional annotations for 36 of them (Fig. S4A). The annotated gene products included DNA metabolism, host cell lysis and structural proteins. The genome also contained 1 tRNA gene. Notably, the phage encodes an Ig domain-containing protein (ACNKOGKT_CDS_0009), associated with interactions between phage capsid and mucin glycoproteins (17). Annotation did not reveal any virulence factors, or gene products associated with integration/excision or AMR. Furthermore, analysis with PhageLeads (18) could not identify any temperate lifestyle markers in the phage genome, classifying fMtkYen801 as suitable for therapeutic applications.

Taxmyphage was unable to classify the phage at genus / species level, suggesting that the phage represents a novel species and genus within the class *Caudoviricetes*.

VIRIDIC analysis of fMtkYen801 and other closely related phages available in GenBank database, showed that query sequence shares highest similarity of 96.2% with *Yersinia* phage fHe-Yen8-01 (GenBank accession number OR620963.1) (Fig. S4B). Viral proteomic tree, constructed based on the genome-wide sequence similarities, identified the host group within the phylum *Pseudomonadota*, confirming experimental data from the host range evaluation, however, virus family could not be detected (Fig. S5A). Phylogenetic analysis using VipTreeGen demonstrated a close evolutionary relationship between fMtkYen801, fHe-Yen8-01, *Escherichia* phage ECML-606-1 (OL631484.1), and *Enterobacter* phage phiT5282EW (PQ352032.1), all of which represents class *Caudoviricetes* (Fig. S5B).

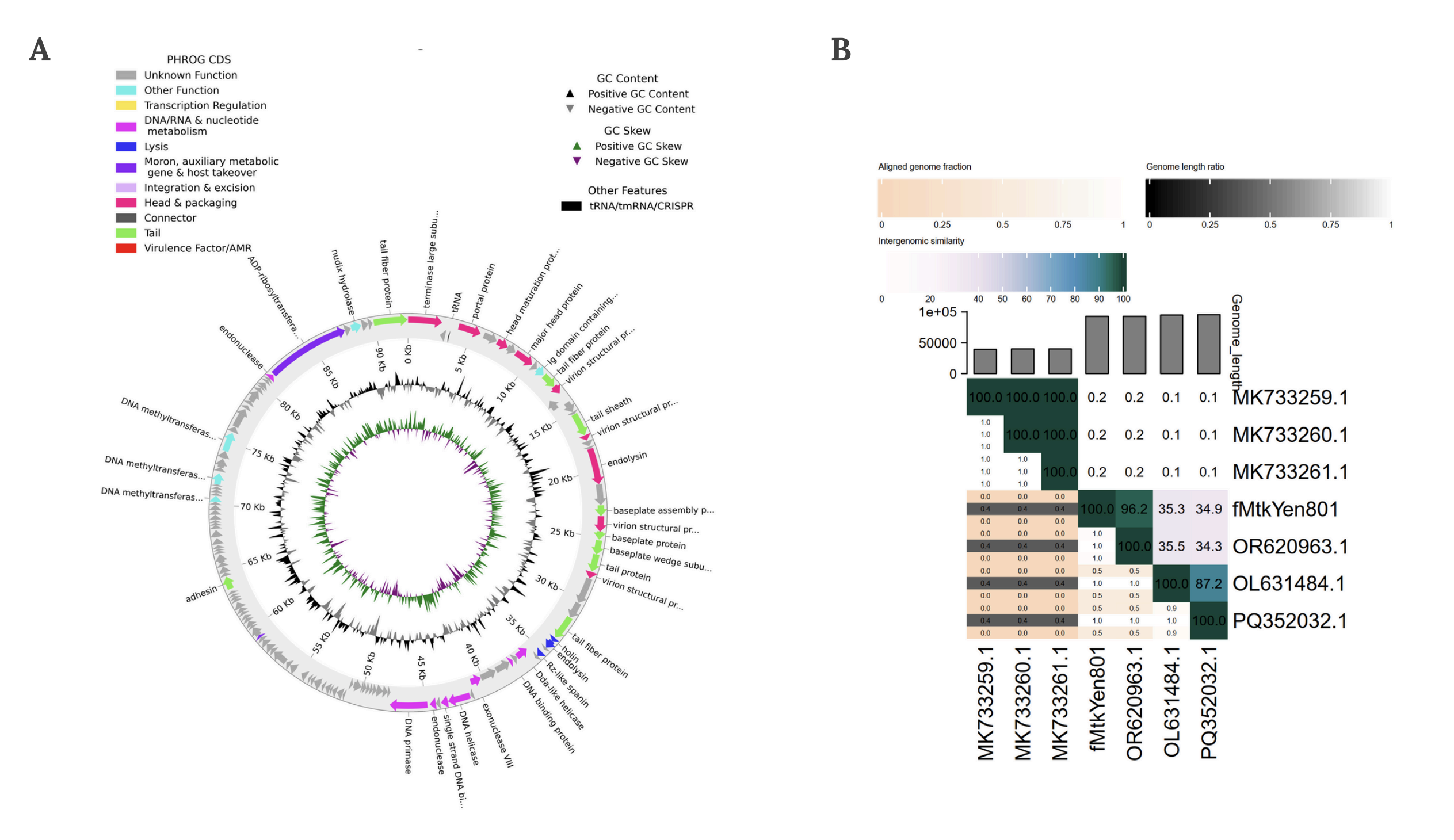

Fig. S4. Genomic analysis of phage fMtkYen801. (A) Genomic map and functional annotation generated using Pharokka. Key structural and functional genes including those involved in DNA metabolism, packaging, or host lysis, are color-coded by functional category (B) Average Nucleotide Identity (ANI) score matrix comparing fMtkYen801 and its closest relative phage genomes, calculated using VIRIDIC

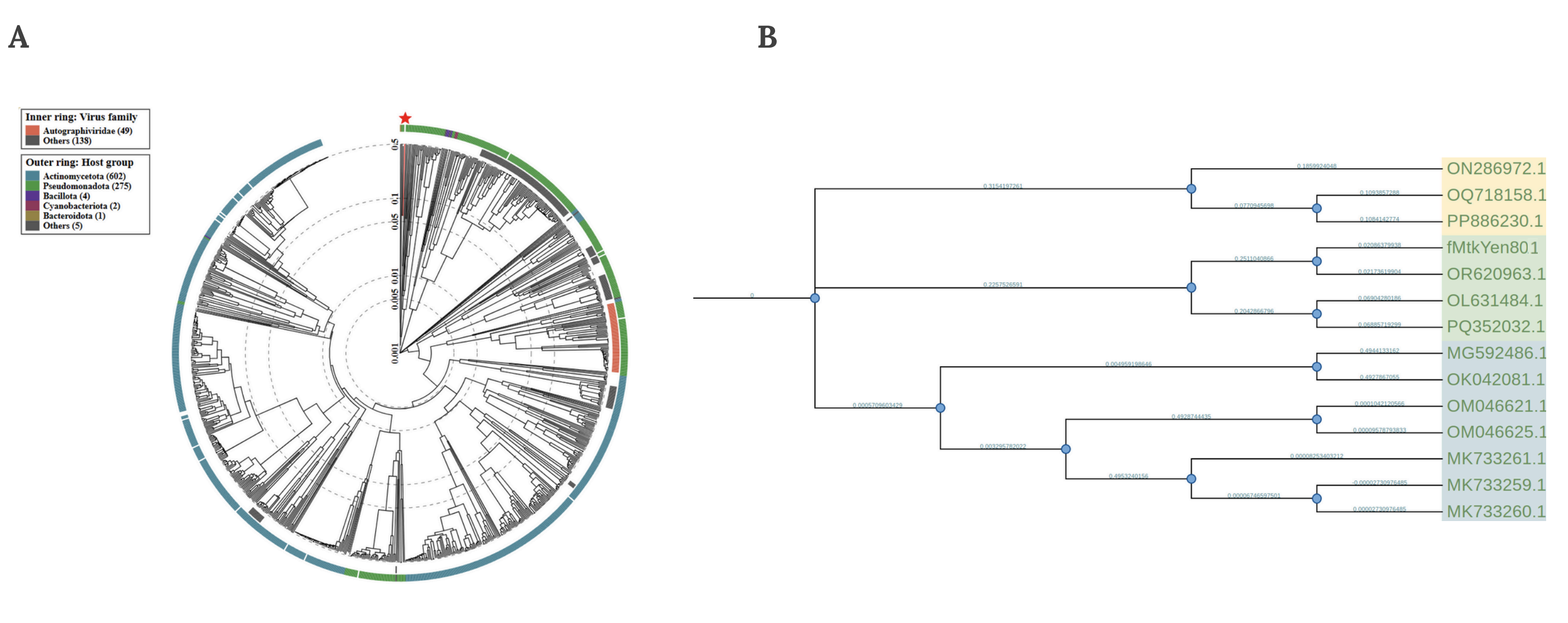

Fig. S5. Phylogenetic analysis of phage fMtkYen801. (A) Viral proteomic (VIP) tree illustrating the evolutionary relationships between fMtkYen801 and closely associated phages (B) Phylogenetic analysis of fMtkYen801 and related *Caudoviricetes* phages based on genome-wide sequence similarity calculated by tBLASTx.

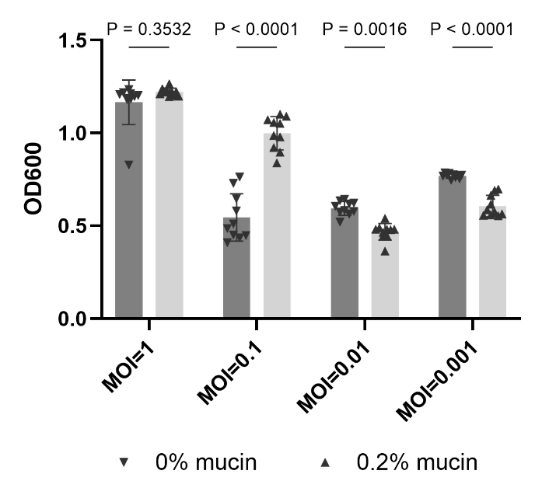

Fig. S6. Abundance of surviving bacterial cells under mucin-enriched (0.2%, w/v) or standard nutrient conditions, determined from optical density (OD_595_) measurements after 60 h time-kill assay with phage fMtkYen801. The error bars represent the mean ± SEM (n = 10 technical replicates per condition). Statistical significance was calculated using 2way ANOVA with Šídák's post hoc test.

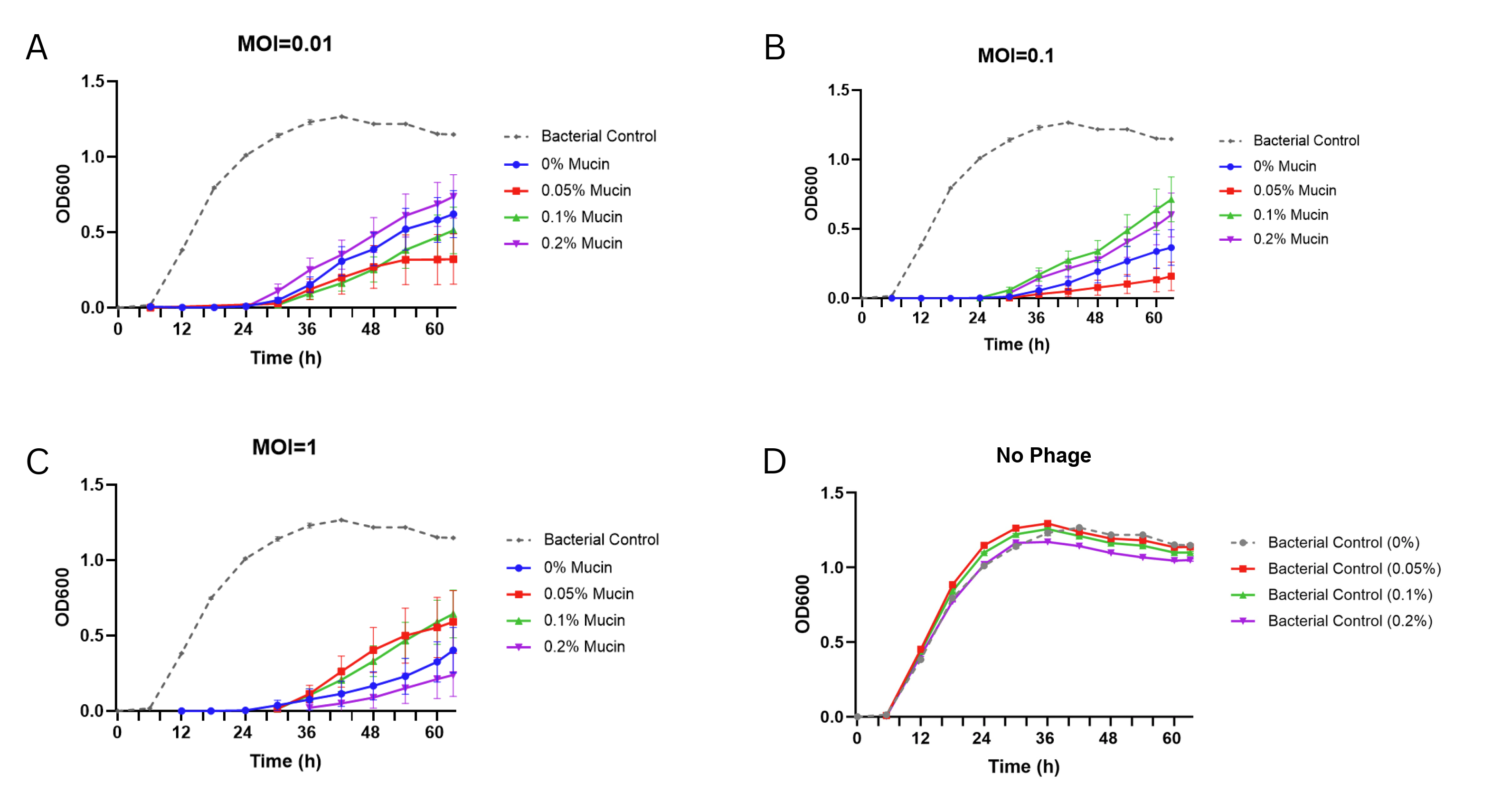

Fig. S7. *Yersinia enterocolitica* O:3 growth curves under gradient of MOI and mucin concentrations, with and without phage fMtkYen3-01 infection. (A) Phage-to-bacteria ratio (MOI) of 0.01 (B) MOI of 0.1 (C) MOI of 1 (D) Bacterial control (no phage). Growth was monitored by OD600 for 60 h, at 25°C. Data are shown as mean ± SEM (n = 10). Statistical significance was calculated using 2way ANOVA with Tukey’s post hoc test.

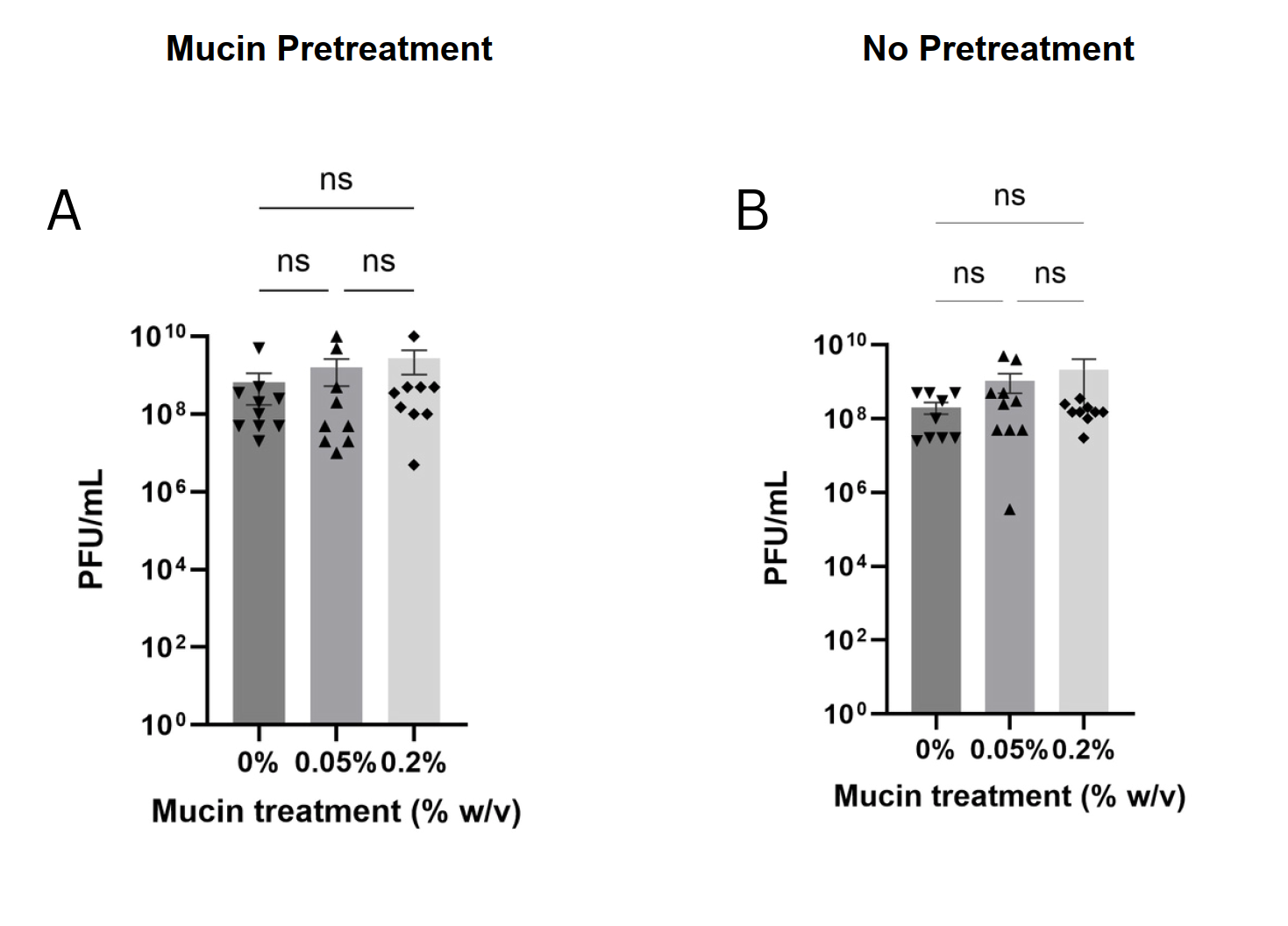

Fig. S8. Phage fMtkYen3-01 abundance after 60 h incubation with *Y. enterocolitica* O:3. (A) Phage titers (PFU/mL) in mucin-pretreated bacterial cultures. (B) Phage replication in bacterial cultures without mucosal pretreatment. Data is shown as mean ±  standard error, n = 10 technical replicates per condition (*p* > 0.05). Statistical significance was calculated using one-way ANOVA followed by Tukey’s post hoc test.

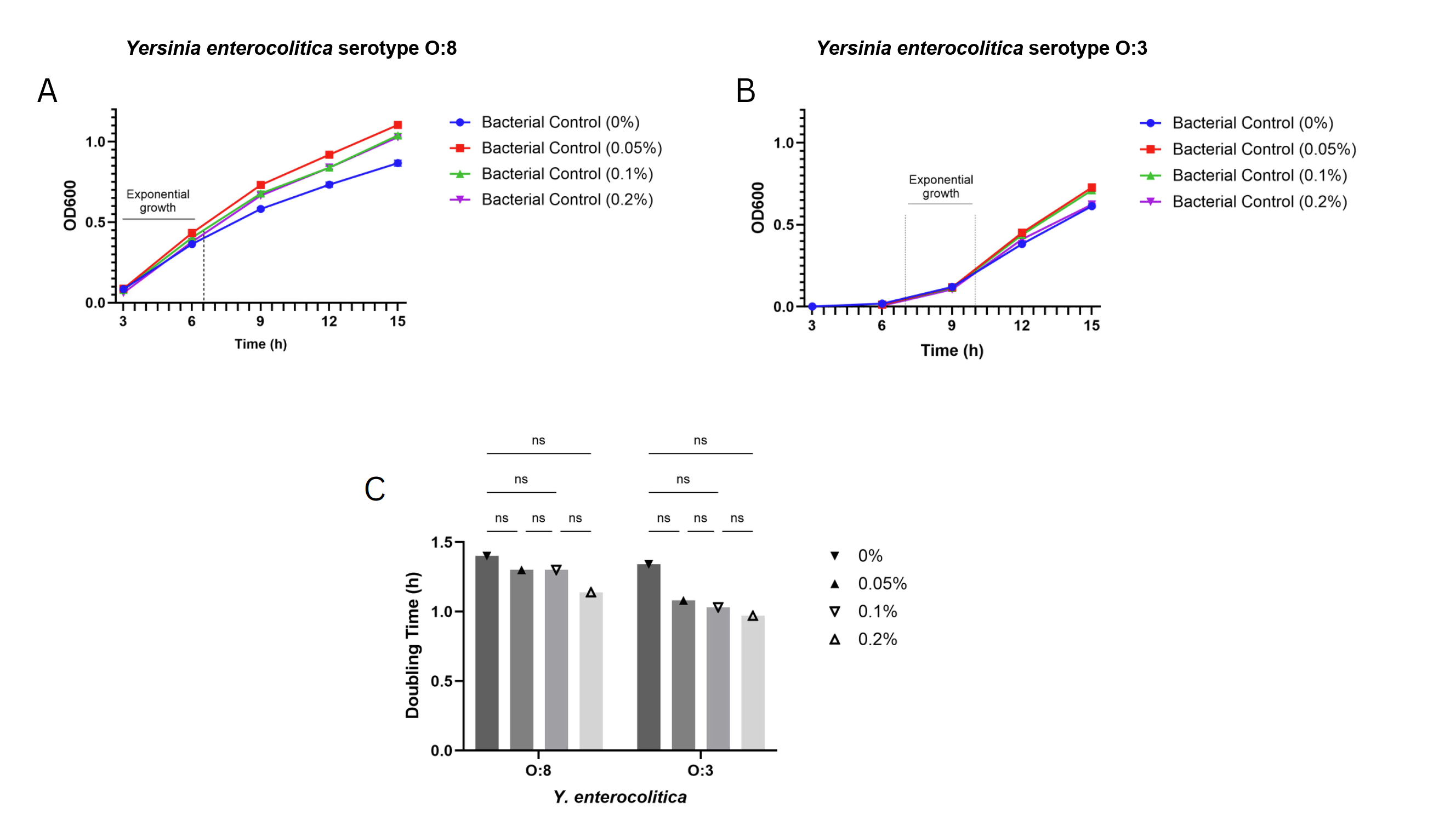

Fig. S9. The effect of mucin supplementation on bacterial doubling time in *Y. enterocolitica* O:8 and *Y. enterocolitica* O:3. (A-B) Growth rates calculated by linear regression of ln-transformed OD_600_ values during the exponential growth phase. (C) Doubling times calculated as ln(2)/μ, where μ is the slope of ln(OD_600_) versus time. Data are shown as mean ± SEM (n = 10 technical replicates) for panels A-B. Statistical significance was assessed using two-way ANOVA with Tukey’s multiple comparison test.

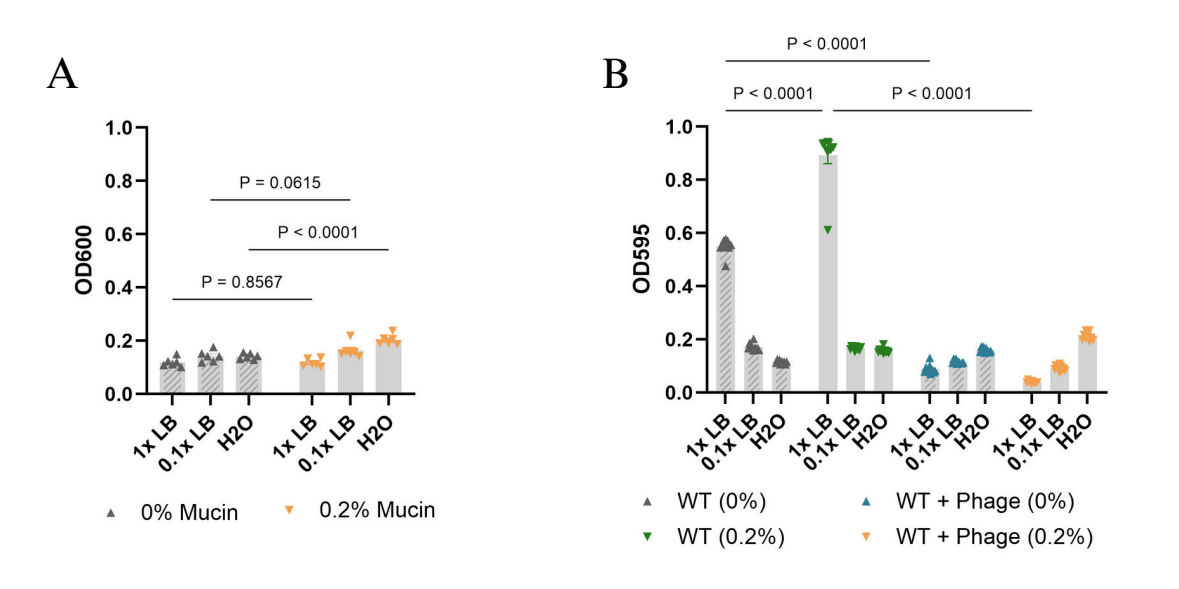

Fig. S10. Control measurements for crystal violet (CV) biofilm assays (Fig. 6). (A) Background CV signal in wells containing sterile medium with or without mucin enrichment (n = 6). (B) Planktonic bacterial cell density (OD_595_) after 48 h static incubation, in the presence or absence of mucin and phage (n = 10). Error bars represent ± SEM. Statistical significance was assessed using two-way ANOVA followed by Tukey’s post hoc test.

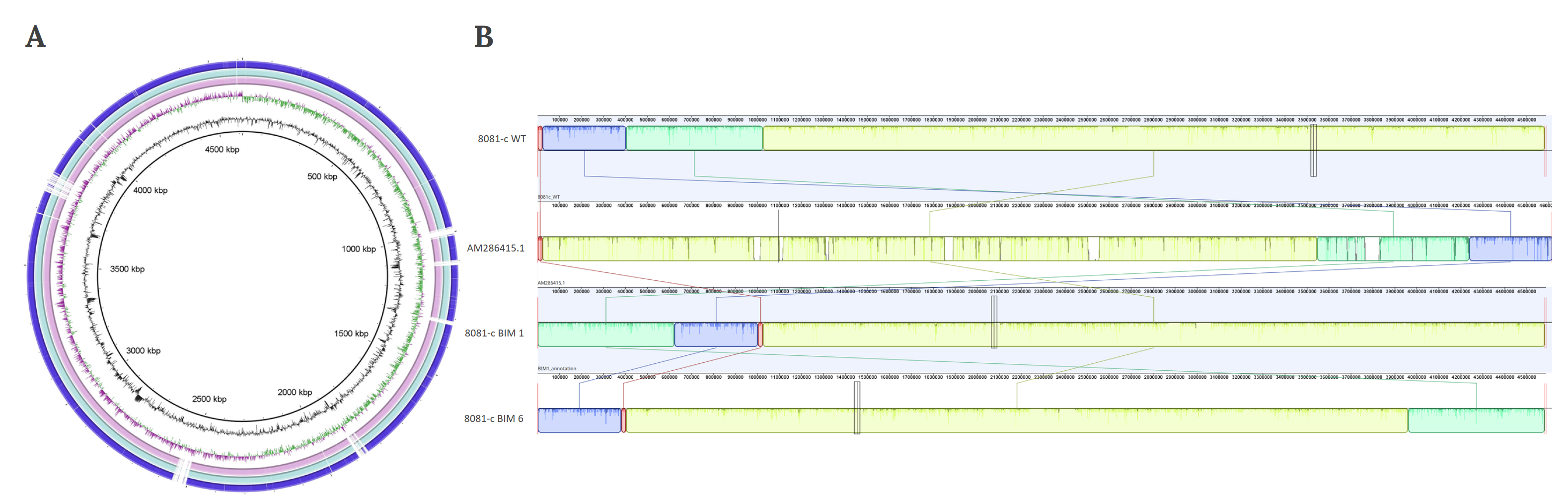

Fig. S11. Comparative genomic analysis of *Y. enterocolitica* 8081c wild-type (WT), its ancestral strain *Y. enterocolitica* 8081, and Bacteriophage Insensitive Mutants (BIMs) (A) Genome comparison visualized using Blast Ring Image Generator (BRIG). The central ring represents the reference genome (*Y. enterocolitica* 8081), while the outer three rings correspond to *Y. enterocolitica* 8081-c WT, BIM 1 from mucin treatment, and BIM 6 from standard nutrient conditions, respectively. (B) Whole genome alignment performed using ProgressiveMauve. Colored collinear blocks (LCBs) show homologous sequences identified between the compared strains. All LCBs are collinear, suggesting structural conservation without major rearrangements between WT and BIM genomes.

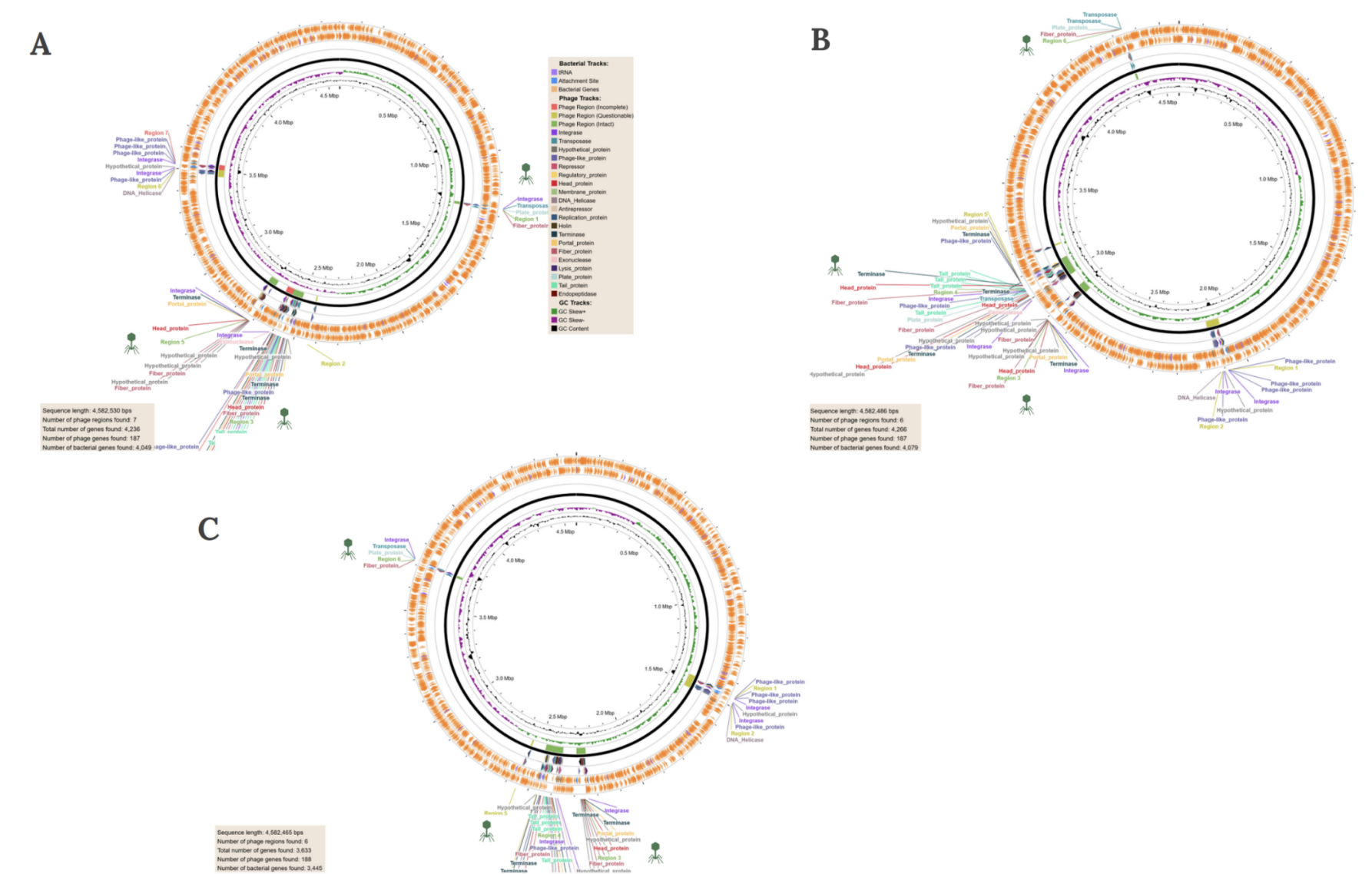

Fig. S12. Circular genome visualized using PHASTEST, highlighting identified prophage regions. Prophages annotated as intact (score > 90) are colored in green across the genome in each panel. Panels show: (A) *Y. enterocolitica* 8081-c WT (B) 8081-c BIM 1 obtained under mucin treatment group, and (C) 8081-c BIM 6 obtained under standard nutrient conditions.

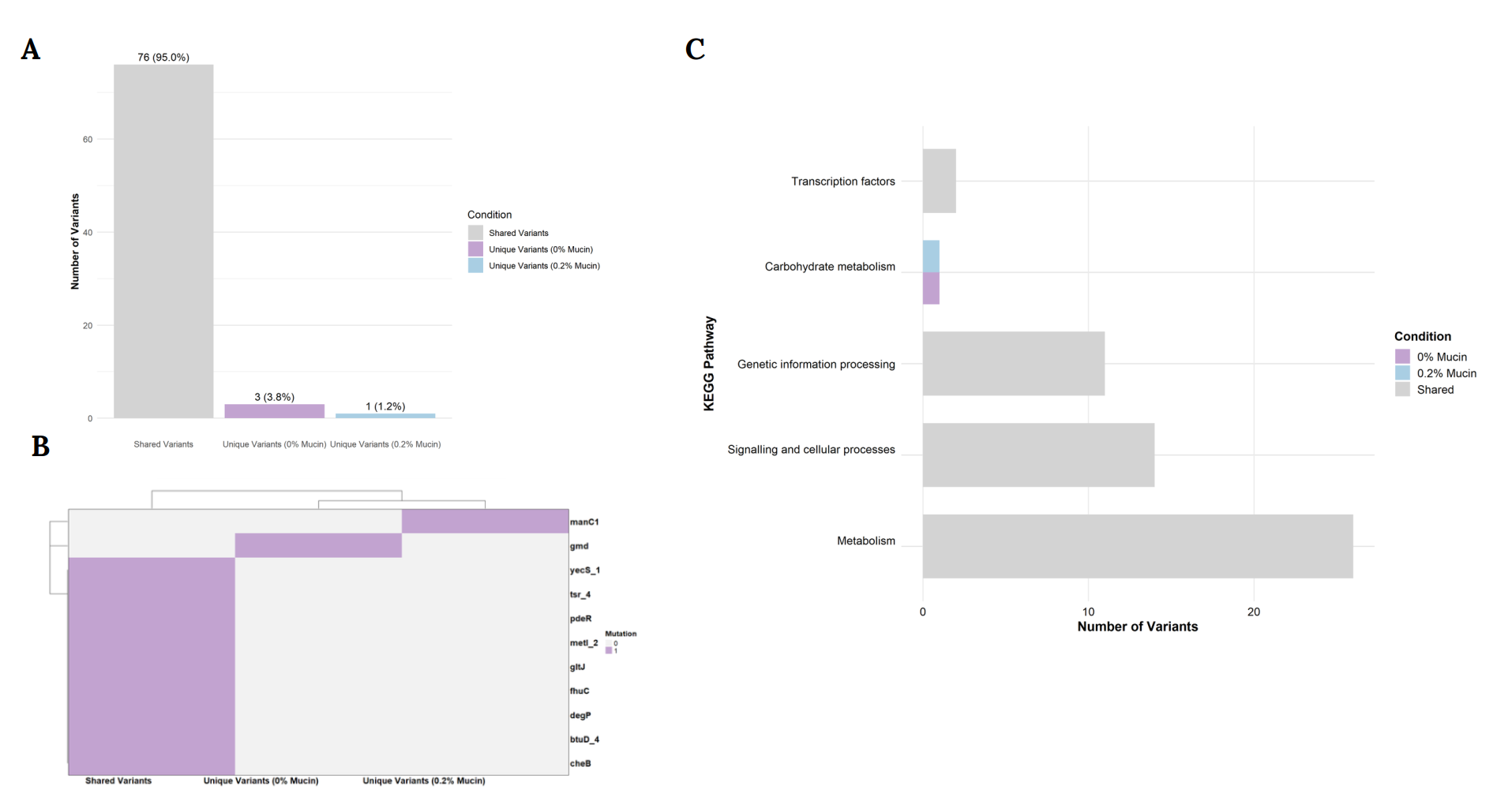

Fig. S13. Variant distribution and gene pathway analysis in Bacteriophage Insensitive Mutants (BIMs) with or without mucin treatment. (A) Percentage distribution of genetic variants, shared by or unique to 0% or 0.2% mucin treatment groups (B) Heatmap visualization of mutations in genes responsible for metabolism, virulence, quorum sensing, phage and antimicrobial resistance by treatment groups, generated using ComplexHeatmap in R (C) Gene pathways affected by phage resistance in each treatment group (0% or 0.2% mucin, w/v), visualized using ggplot2.

**References:**

1.	Gomez-Raya-Vilanova MV, Leskinen K, Bhattacharjee A, Virta P, Rosenqvist P, Smith JLR, et al. The DNA polymerase of bacteriophage YerA41 replicates its T-modified DNA in a primer-independent manner. Nucleic Acids Res. 2022 Apr 22;50(7):3985–97.

2.	Leskinen K, Tuomala H, Wicklund A, Horsma-Heikkinen J, Kuusela P, Skurnik M, et al. Characterization of vB_SauM-fRuSau02, a Twort-Like Bacteriophage Isolated from a Therapeutic Phage Cocktail. Viruses. 2017 Sep;9(9):258.

3.	Tritt A, Eisen JA, Facciotti MT, Darling AE. An Integrated Pipeline for de Novo Assembly of Microbial Genomes. PLOS ONE. 2012 Sep 13;7(9):e42304.

4.	PhageTerm: a tool for fast and accurate determination of phage termini and packaging mechanism using next-generation sequencing data | Scientific Reports [Internet]. [cited 2025 Sep 30]. Available from: https://www.nature.com/articles/s41598-017-07910-5

5.	Bouras G, Nepal R, Houtak G, Psaltis AJ, Wormald PJ, Vreugde S. Pharokka: a fast scalable bacteriophage annotation tool. Bioinformatics. 2023 Jan 1;39(1):btac776.

6.	Millard A, Denise R, Lestido M, Thomas MT, Webster D, Turner D, et al. taxMyPhage: Automated Taxonomy of dsDNA Phage Genomes at the Genus and Species Level. Phage (New Rochelle). 2025 Mar;6(1):5–11.

7.	Moraru C, Varsani A, Kropinski AM. VIRIDIC—A Novel Tool to Calculate the Intergenomic Similarities of Prokaryote-Infecting Viruses. Viruses. 2020 Nov;12(11):1268.

8.	Nishimura Y, Yoshida T, Kuronishi M, Uehara H, Ogata H, Goto S. ViPTree: the viral proteomic tree server. Bioinformatics. 2017 Aug 1;33(15):2379–80.

9.	Letunic I, Bork P. Interactive Tree of Life (iTOL) v6: recent updates to the phylogenetic tree display and annotation tool. Nucleic Acids Res. 2024 Jul 5;52(W1):W78–82.

10.	Goladze S, Patpatia S, Tuomala H, Ylänne M, Gachechiladze N, de Oliveira Patricio D, et al. Isolation and characterization of Yersinia phage fMtkYen3-01. Arch Virol. 2024 Oct 19;169(11):226.

11.	Skurnik M, Alkalay-Oren S, Boon M, Clokie M, Sicheritz-Pontén T, Dąbrowska K, et al. Phage therapy. Nat Rev Methods Primers. 2025 Feb 13;5(1):9.

12.	Patpatia S, Schaedig E, Dirks A, Paasonen L, Skurnik M, Kiljunen S. Rapid hydrogel-based phage susceptibility test for pathogenic bacteria. Frontiers in Cellular and Infection Microbiology [Internet]. 2022 [cited 2023 Apr 2];12. Available from: https://www.frontiersin.org/articles/10.3389/fcimb.2022.1032052

13.	Jun JW, Park SC, Wicklund A, Skurnik M. Bacteriophages reduce Yersinia enterocolitica contamination of food and kitchenware. International Journal of Food Microbiology. 2018 Apr 20;271:33–47.

14.	Mirzaei MK, Nilsson AS. Isolation of Phages for Phage Therapy: A Comparison of Spot Tests and Efficiency of Plating Analyses for Determination of Host Range and Efficacy. PLOS ONE. 2015 Mar 11;10(3):e0118557.

15.	Bengoechea JA, Najdenski H, Skurnik M. Lipopolysaccharide O antigen status of Yersinia enterocolitica O:8 is essential for virulence and absence of O antigen affects the expression of other Yersinia virulence factors. Molecular Microbiology. 2004 Apr 1;52(2):451–69.

16.	Colom J, Cano-Sarabia M, Otero J, Aríñez-Soriano J, Cortés P, Maspoch D, et al. Microencapsulation with alginate/CaCO3: A strategy for improved phage therapy. Sci Rep. 2017 Jan 25;7(1):41441.

17.	Barr JJ, Auro R, Furlan M, Whiteson KL, Erb ML, Pogliano J, et al. Bacteriophage adhering to mucus provide a non–host-derived immunity. Proceedings of the National Academy of Sciences. 2013 Jun 25;110(26):10771–6.

18.	Yukgehnaish K, Rajandas H, Parimannan S, Manickam R, Marimuthu K, Petersen B, et al. PhageLeads: Rapid Assessment of Phage Therapeutic Suitability Using an Ensemble Machine Learning Approach. Viruses. 2022 Feb;14(2):342.
